# Membrane phosphoinositides stabilize GPCR-arrestin complexes and provide temporal control of complex assembly and dynamics

**DOI:** 10.1101/2021.10.09.463790

**Authors:** John Janetzko, Ryoji Kise, Benjamin Barsi-Rhyne, Dirk H. Siepe, Franziska M. Heydenreich, Matthieu Masureel, Kouki Kawakami, K. Christopher Garcia, Mark von Zastrow, Asuka Inoue, Brian K. Kobilka

## Abstract

Binding of arrestin to phosphorylated G protein-coupled receptors (GPCRs) is crucial for modulating signaling. Once internalized some GPCRs may complex with arrestin, while others interact transiently; this difference affects receptor signaling and recycling. Cell-based and in vitro biophysical assays reveal the role of membrane phosphoinositides (PIPs) in arrestin recruitment and GPCR-arrestin complex dynamics. We find that GPCRs broadly stratify into two groups, one requiring PIP-binding for arrestin recruitment and one that does not. Plasma membrane PIPs potentiate an active conformation of arrestin and stabilize GPCR-arrestin complexes by promoting a receptor core-engaged state of the complex. As allosteric modulators of GPCR-arrestin complex dynamics, membrane PIPs allow for additional conformational diversity beyond that imposed by GPCR phosphorylation alone. The dependance on membrane PIPs provides a mechanism for arrestin release from transiently associated GPCRs, allowing their rapid recycling, while explaining how stably associated GPCRs are able to engage G proteins at endosomes.

## Introduction

G protein-coupled receptor (GPCR) activation and deactivation are tightly regulated, allowing them to achieve robust signaling. GPCR deactivation is a complex multi-step process often divided into an acute and a prolonged phase (Rajagopal and Shenoy, 2018). In addition to promoting G protein engagement, agonist stimulation leads to the recruitment of GPCR kinases (GRKs), which phosphorylate the receptor and trigger recruitment of arrestins (Komolov and Benovic, 2018). Arrestin first blocks further G protein engagement, resulting in an acute phase of desensitization, but also mediates the trafficking of activated receptors to clathrin-coated structures (CCSs) and their internalization. Once internalized, receptors can experience markedly different fates, with some being rapidly recycled to the plasma membrane, while others are retained in intracellular compartments, or directed to lysosomes and degraded (Hanyaloglu and von Zastrow, 2008). In recent years, the discovery that GPCRs can signal from intracellular compartments has led to a re-framing of GPCR signaling to include not only temporal regulation, but also differences that result from spatially distinct receptor populations (Irannejad et al., 2013; Irannejad et al., 2015; Lobingier and von Zastrow, 2019).

There are four human arrestins; arrestins 1 and 4 are dedicated to the visual system, and arrestins 2 and 3, also known as β-arrestin 1 (βarr1) and β-arrestin 2 (βarr2), respectively, are ubiquitously expressed throughout the other tissues. Remarkably, these two β-arrestins are responsible for recognition and desensitization of hundreds of GPCRs. Though most GPCRs recruit arrestin, the nature and duration of this interaction can differ between receptors. Historically GPCRs have been classified as either a “class A” receptor, which interacts transiently with arrestin, or “class B” receptors, which interact more stably with arrestin and co-localize with arrestin in endosomes (Oakley et al., 2001; Oakley et al., 2000; Zhang et al., 1999). This distinction is importantly different from that of family A (rhodopsin-like) and family B (secretin-like) GPCRs. Whether a GPCR interacted transiently or stably with arrestin appears to correlate with rates of re- sensitization, with class A receptors re-sensitizing more rapidly than class B receptors (Oakley et al., 1999). Stable association of arrestin to “class B” GPCRs correlates with the presence of particular phosphorylation site clusters (Oakley et al., 2001); however, it has remained unknown what event precipitates the dissociation of β-arrestins from “class A” receptors to allow their dephosphorylation and recycling.

Early structural studies into GPCR-arrestin complexes suggested that arrestin binds to a GPCR either through only the phosphorylated C-terminus (called tail-engaged), or through both the phosphorylated C-terminus and the transmembrane core of the GPCR (called core-engaged) (Latorraca et al., 2018; Shukla et al., 2014; Staus et al., 2018). However, it remains unclear what determines the equilibrium between these states. While core-engagement is necessary for receptor desensitization (Kumari et al., 2017), it is not required for internalization (Cahill et al., 2017). If, however, complexes can shift from core-engaged to tail-engaged in endosomes it would allow for G proteins to access the receptor core while remaining bound to arrestin. These so-called “megaplex” assemblies (Nguyen et al., 2019; Thomsen et al., 2016) have been implicated in the sustained cAMP signaling produced by endosomal populations of V2R and PTH1R (Feinstein et al., 2013; Ferrandon et al., 2009), both of which stably associate with β-arrestins.

At a molecular level, the prevailing model for arrestin activation, and thus recruitment to an active and phosphorylated GPCR, involves displacement of the auto-inhibitory C-terminus of arrestin by the GPCR phosphorylated C-terminus (or in some cases an intracellular loop). Once the arrestin C-terminus is displaced, additional structural rearrangements occur that allow for arrestin to engage a GPCR (Sente et al., 2018), including insertion of the arrestin finger loop into a cavity formed by the cytoplasmic ends of transmembrane segments. However, arrestin activation functions for more than just GPCR engagement. In its active form, arrestin is able to engage multiple signaling proteins, including JNK3, ERK1/2, p38 (Song et al., 2009) and Src (Pakharukova et al., 2020). It has been suggested that distinct arrestin conformations, which can arise from different inputs (i.e., receptor phosphorylation pattern) may favor interaction with a subset of these signaling partners and affect signaling outcomes downstream of arrestin (Chen et al., 2018; Latorraca et al., 2020). Recently the model for arrestin activation, which suggests a 1:1 interaction, has been challenged by the finding that some “class A” receptors cause the accumulation of super-stoichiometric quantities of arrestin in clathrin-coated structures (CCSs), suggesting an ability to persist at the membrane without an associated GPCR (Eichel et al., 2018; Nuber et al., 2016). It was speculated that an association with PIP_2_ was responsible for retaining β-arrestins in clathrin coated structures (Eichel et al., 2018); however, based on the established mechanism for arrestin activation it is unclear how this would be possible, or how arrestin absent an associated GPCR promoted MAPK signaling from CCSs (Eichel et al., 2016).

Components of the endocytic machinery such as AP2 (Kadlecova et al., 2017), and β-arrestins (Gaidarov et al., 1999) have been shown to bind to PIPs. These signaling lipids serve critical functions defining the identity of lipid compartments and acting as coincidence markers for protein-protein recognition and trafficking to occur only in the appropriate subcellular context (De Matteis and Godi, 2004; Di Paolo and De Camilli, 2006). While several studies have investigated the interactions of soluble inositol phosphates with both visual and non-visual arrestins (Chen et al., 2017; Chen et al., 2021; Milano et al., 2006; Zhuang et al., 2010),there has been only one where the role of membrane PIPs was explored (Gaidarov et al., 1999). Importantly, this work suggested that plasma membrane PIPs, such as PI(4,5)P2 and PI(3,4,5)P3, hereafter PIP_2_ and PIP_3_, respectively, may function to stabilize GPCR-β-arrestin complexes as they traffic to CCSs.

Recent structural studies showing PIP_2_ bound at the interface between the neurotensin type I receptor (NTSR1) and βarr1 (Huang et al., 2020) prompted us to ask the question: what role do PIPs serve in mediating GPCR-β-arrestin complex assembly? Here we show that some GPCRs require PIP binding for β-arrestin recruitment, provided they engage β-arrestins transiently. Furthermore, we show that the requirement of PIPs depends on specific receptor phosphorylation sites. Using *in vitro* biochemical and biophysical assays, we demonstrate that phosphoinositide binding contributes to the stability of the GPCR-β-arrestin complex, where it promotes the core-engaged state. We also find that PIPs alone promote a partially activated state of arrestin, providing an explanation for how arrestin can persist in CCSs once dissociated from a GPCR. Together, these results explain a) how receptors that transiently associate with β-arrestin recruit and dissociate β-arrestin in a spatiotemporally resolved manner, and b) how strongly coupled receptors maintain a stable association with arrestin in subcellular compartments yet allow for further G protein engagement from subcellular structures.

## Results and Discussion

### Arrestin PIP-binding is important for desensitization of endogenous β2AR

The PIP-binding-deficient mutant of βarr2 (K233Q/R237Q/K251Q, henceforth 3Q, also used to denote mutation of the homologous residues in K232Q/R236Q/K250Q in βarr1) was previously found to be impaired for internalization of β2AR (Gaidarov et al., 1999), with βarr2 (3Q) failing to traffic to CCSs, though still being recruited from the cytoplasm to the plasma membrane, albeit to a lesser extent than wild-type (WT) (Eichel et al., 2018). As such, we wondered how this behavior affects β2AR signaling and specifically whether βarr2 (3Q) is capable of desensitizing β2AR at the plasma membrane. Using a FRET-based cAMP sensor (Tewson et al., 2016), we monitored cAMP production in real-time in HEK293 cells lacking both β-arrestins, and endogenously expressing the β2AR (O’Hayre et al., 2017). In the absence of exogenously expressed βarr2, isoproterenol (iso) stimulation, via endogenous β2AR, led to a sustained cAMP response, while expression of βarr2 led to desensitization. However, expression of the 3Q βarr2 mutant resulted in significantly less desensitization over 30 minutes (Figure S1A); furthermore, this difference was observed in two independent β-arrestin-deficient cell lines (Luttrell et al., 2018; O’Hayre et al., 2017). This suggests that the PIP-binding function of β-arrestins plays an important functional role, in not only internalization (Gaidarov et al., 1999), but also receptor desensitization, and does so under conditions of endogenous GPCR expression.

### GPCRs stratify into two groups in their dependence on PIP-binding for arrestin recruitment

That the βarr2 3Q mutant is impaired for recruitment to β2AR, but seemingly not for the chimeric receptor β2AR-V2C, which bears the C-terminus of the vasopressin V2 receptor (Eichel et al., 2018), suggested that GPCRs may differ in their dependence on β-arrestin PIP-binding for recruitment. To investigate a wide range of GPCRs we used a cell-based NanoBiT assay (Dixon et al., 2016), wherein a plasma membrane localization sequence (CAAX) is fused to the large subunit of a modified NanoLuc luciferase (LgBiT). Recruitment of either βarr1 or βarr2, which bear an N-terminal complementary small subunit of NanoLuc (SmBiT), can be monitored by luminescence changes (Figure 1A). We selected a set of 22 representative GPCRs (Supplementary Data Table 1), co-expressed the sensors with each receptor of interest in HEK293 cells, and compared the recruitment of WT β-arrestin to that of the corresponding 3Q β- arrestin mutant upon agonist stimulation (Figure 1B, top, Supplementary Data Table 1). We used luminescence fold-change measured over the range of 10-15 minutes post-agonist stimulation for our end-point values and fit the resulting data to generate concentration response curves and extract a recruitment amplitude for each receptor-arrestin pair (see methods) (Figure 1B, bottom, Supplementary Data Figure 1A-B). We then compared the recruitment of WT and 3Q arrestin using a metric that represented the relative sensitivity of the receptor to loss of arrestin-PIP binding capacity, we termed the loss of function (LOF) index (see methods). Receptors with a low LOF value recruit WT and 3Q β-arrestins to the plasma membrane similarly, and are deemed PIP-independent, while receptors with a high LOF value show greatly diminished recruitment of 3Q β-arrestin and are deemed PIP-dependent (Figure 1C). Both WT and 3Q forms of βarr1 and βarr2 express similarly (Figure S1B, Supplementary Data Figure 2).

**Figure 1.**
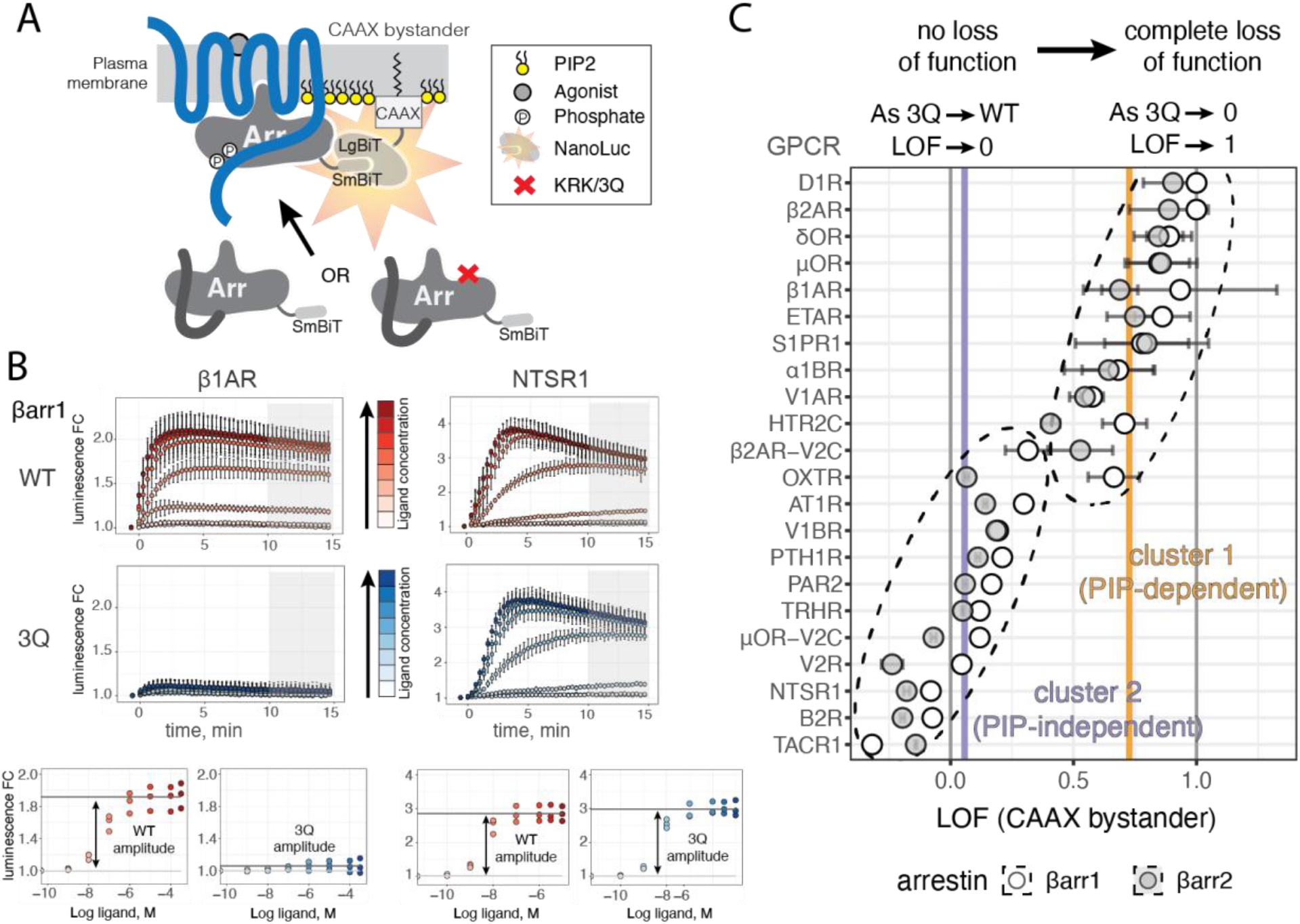
Arrestin phosphoinositide binding is required for recruitment to some GPCRs A) cartoon depicting NanoBiT assay for measuring arrestin plasma membrane recruitment upon agonist stimulation. Upon complementation SmBiT and LgBiT form a functional NanoLuc luciferase. In key, “Phosphate” denotes phosphorylated Ser/Thr residues and “X” denotes KRK to 3Q mutant of β-arrestin. B) Two representative GPCRs, β1AR and NTSR1 illustrate data obtained for β- arrestin recruitment by NanoBiT assay shown in panel A. Data were collected over time after agonist addition (t=0 min), and values are shown as luminescence fold-change (over vehicle treatment) ± standard deviation (measured as 2 technical replicates for each of *n*=3 independent experiments). Colors denote concentrations of agonist used for stimulation. Agonists used were isoproterenol for β1AR and neurotensin for NTSR1. Grey boxes mark the time region (10-15 minutes post agonist addition) over which luminescence is integrated, for each concentration of agonist, to produce concentration response curves (bottom). WT and 3Q amplitudes were determined as the difference of fitted pre- and post-transition plateaus. C) Plot of LOF values for panel of tested GPCRs. Points represent LOF value obtained as ratio of WT and 3Q recruitment, and error bars reflect error in LOF derived from standard errors of fits (see methods). Dashed ellipses denote clusters obtained from k means clustering of data (see methods); AT1R is in cluster 2 for both βarr1/2, while β2AR-V2C is split with βarr1 (cluster 2) and βarr2 (cluster 1). Vertical grey lines denote LOF = 0 and LOF =1; vertical purple and orange lines reflect the centers of the respective clusters from k means and correspond to LOF = 0.06 and LOF = 0.73, respectively.

**Figure 2.**
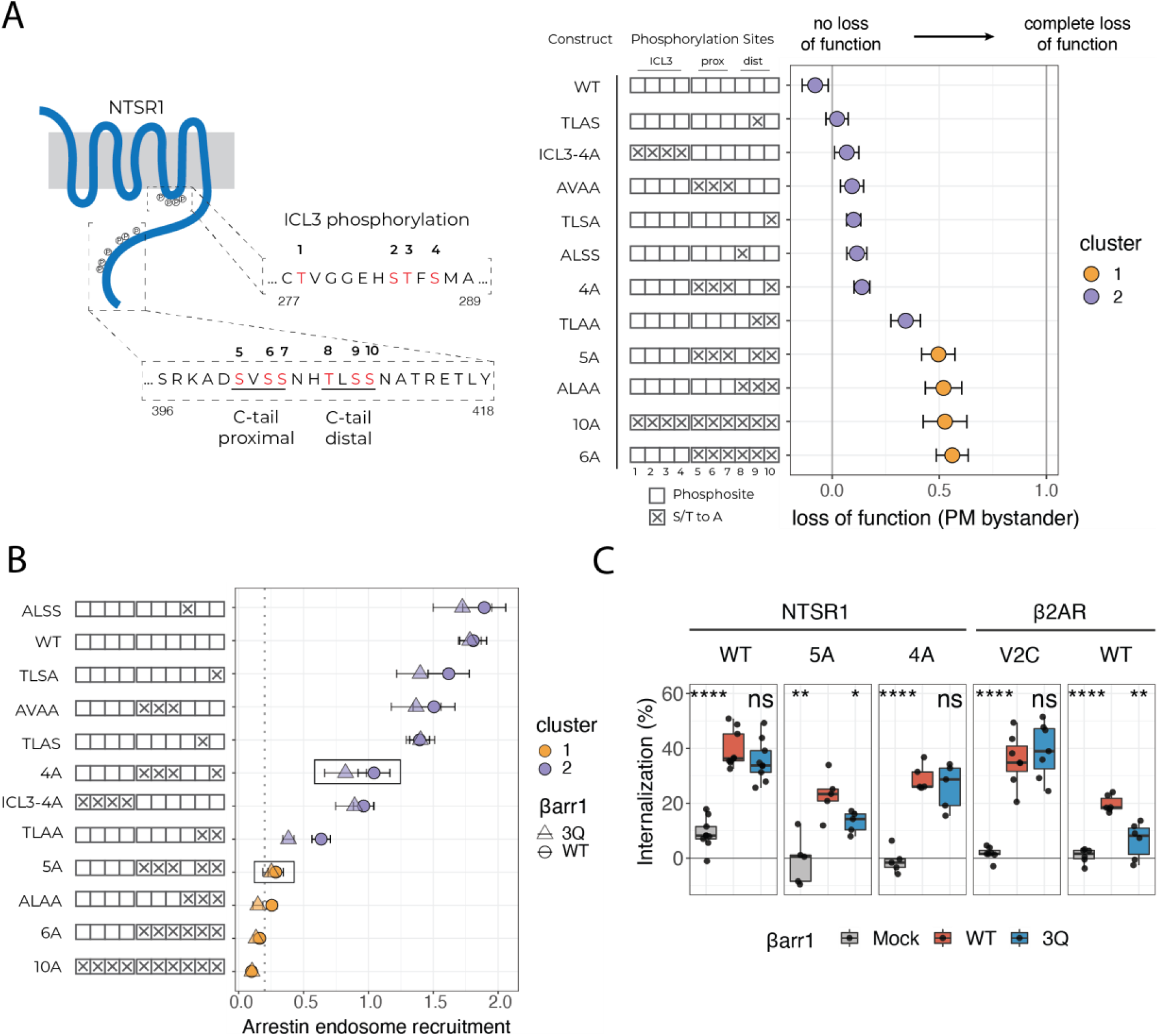
Receptor phosphorylation patterns govern PIP-dependence for arrestin recruitment. A) Left, schematic of human NTSR1 showing motifs in receptor ICL3 and C-terminus that are subject to phosphorylation. Phosphorylation sites examined in this study are shown in red and numbered 1-10 (above). Residue numbers corresponding to the region of human NTSR1 are listed at the start and end of the shown sequences. Construct key shows possible phosphosites as empty boxes, which when mutated to alanine are filled with an “X”. Plasma membrane recruitment of arrestin upon stimulation of cells expressing different NTSR1 constructs, measured using the NanoBiT assay described in Figure 1. Right, points represent LOF value obtained as ratio of WT and 3Q recruitment, and error bars represent standard error of fits (see methods). Points are colored based on cluster designation obtained from k means clustering of all receptor-arrestin recruitment data. B) Translocation of βarr1 to endosomes upon stimulation of cells expressing different NTSR1 constructs, measured using an endosome bystander NanoBiT assay, as described in Figure S1. Points represent recruitment (fold chance over basal upon stimulation) for WT and 3Q recruitment, denoted by circles and triangles, respectively. Points are based on data from n=3 biological experiments. Error bars represent standard error of fit used to determine recruitment. Points are colored based on the cluster assignment of that mutant. C) Internalization, measured by loss of cell-surface receptors upon agonist stimulation, for Δβarr1/2 cells expressing NTSR1 or β2AR constructs and transfected with arrestin constructs indicated. Values represent independent experiments (*n* = 5-10). Internalization by 3Q βarr1 and mock were compared to WT using a two-tailed paired t-test. ns: p > 0.05; *: p ≤ 0.05; **: p ≤ 0.01; ***: p ≤ 0.001; ****: p ≤ 0.0001.

Though receptors spanned a continuum of LOF values, they seemed to cluster into two groups near the ends of the scale. To examine this, we performed k means clustering of plasma membrane recruitment data for all GPCR-β-arrestin pairs (Supplementary Data Figures 1 and 3, *n* = 55 pairs), which suggested that the data is best divided into two clusters (see methods, clusters marked by dotted ellipses in Figure 1C). We found only a weak inverse correlation between the amplitude of WT arrestin recruitment and the degree of LOF observed (Pearson correlation = -0.51; -0.4 when TACR1 and B2R are excluded) (Figure S1C), suggesting that differences in LOF were not due to lower levels of WT recruitment. Cluster 1 was defined by receptors that exhibited a high degree of LOF (center LOF = 0.73) and included GPCRs previously classified as “class A” (Oakley et al., 2000): β2AR, μOR, ETAR, D1R, α1BR. Cluster 2, defined by receptors with a low degree of LOF (center LOF = 0.06), included GPCRs classified as “class B” (Oakley et al., 2000): AT1R, NTSR1, V2R, TRHR, and TACR1. We also tested two chimeric receptors, β2AR-V2C and μOR-V2C (Eichel et al., 2018), both of which showed reduced reliance on β-arrestin PIP-binding capability for plasma membrane recruitment compared to the respective parent receptor. The V1AR, which was previously shown to undergo labile phosphorylation and rapid recycling (Innamorati et al., 1998a; Innamorati et al., 1998b) clusters with the class A receptors in cluster 1, while the V1BR, bearing a closer similarity in its proximal C-terminus to V2R clusters with class B receptors in cluster 2, even though it has been found to only associate transiently with arrestin (Perkovska et al., 2018). In addition, β1AR, S1PR1, and δOR, all three of which have been shown to either recycle rapidly or interact transiently with arrestin (Martinez-Morales et al., 2018; Nakagawa and Asahi, 2013; Trapaidze et al., 2000), were assigned to cluster 1. Other receptors known to co-localize with arrestin at endosomes, including PAR2 (DeFea et al., 2000; Dery et al., 1999; Oakley et al., 2001), B2R (Khoury et al., 2014), and PTH1R (Feinstein et al., 2011) were also classified into cluster 2. Two receptors, OXTR and HTR2C displayed unexpected behavior where βarr1 recruitment was dramatically more sensitive to loss of PIP- binding than βarr2, resulting in these GPCR-β-arrestin pairs being divided between the two clusters. OXTR was previously classified as a “class B” receptor (Oakley et al., 2001); however, these studies only examined βarr2 recruitment. For the serotonin 2C receptor (HTR2C), previous studies showed PIP_2_-depletion did not block recruitment of βarr2 (Toth et al., 2012), and we observed an intermediate LOF value for βarr2 plasma membrane recruitment with HTR2C. Together, these data show that recruitment of β-arrestins is dependent on the PIP-binding capacity of arrestin for some GPCRs, but not others, and that this distinction is consistent with the previous class A/B categorization based on microscopy co-localization studies.

While our use of a plasma membrane localized LgBiT avoids modifying the receptor of interest, we wanted to confirm that plasma membrane recruitment is indeed a reliable proxy for arrestin recruitment to a GPCR of interest. For this, we used a direct NanoBiT assay in which the SmBiT component is fused to the C-terminus of each GPCR of interest, and the N-terminus of arrestin is modified with the LgBiT fragment (Figure S1D, left). We found that recruitment measured by direct complementation largely paralleled recruitment measured using the plasma membrane bystander, with minor exceptions (Supplementary Data Figure 4-5). Further, directly comparing LOF as measured by the plasma membrane bystander to that of the direct complementation showed a strong positive correlation (Pearson correlation = 0.88), suggesting that β-arrestin recruitment measured through the plasma membrane bystander was indeed a faithful metric (Figure S1D, right). The most extreme outlier, HTR2C, showed βarr2 recruitment is PIP-binding independent as measured by the direct recruitment assay, as compared to being partially PIP-binding-dependent when measured using the plasma membrane bystander. As the direct recruitment assay seemed to better match prior findings for the HTR2C (Toth et al., 2012), we wondered why this might be. In addition to HTR2C, α1BR and β1AR also exhibit reduced PIP-binding sensitivity in the direct recruitment assay for βarr2. Curiously, all three of these receptors exhibit some level of Gq coupling (Inoue et al., 2019). We speculate that for cluster 1 receptors, such as HTR2C, which are Gq-coupled, that their dependence on PIP-binding for arrestin recruitment to the plasma membrane may be amplified by local PIP_2_-depletion via phospholipase C upon stimulation.

Given that “class B” receptors co-localize with β-arrestins in endosomes, we wondered whether PIP binding affected this process. We used the FYVE domain of endofin as an endosome bystander (Endo) (Namkung et al., 2016), which we fused to LgBiT (termed endo-Lg) to monitor recruitment of arrestin bearing an N-terminal SmBiT (Figure S1E), as was done for plasma membrane recruitment. Since both βarr1 and βarr2 displayed largely similar behavior in our plasma membrane recruitment assay, we focused on βarr1 for these experiments (Supplementary Data Figure 6A); however, βarr2 recruitment was also examined for a subset of receptors (Supplementary Data Figure 6B). This assay robustly detected endosomal translocation as all receptors known to co-localize with β-arrestin in endosomes did so (Figure S1E). Consistent with the stark difference between βarr1 and βarr2 observed for OXTR (Figure 1C), we observed measurable endosomal association of βarr2, but weak and barely measurable βarr1 endosome recruitment (Supplementary Data Figure 6C). Our results were consistent with prior microscopy-based approaches (Oakley et al., 2000), thereby validating our NanoBiT assay. In contrast, HTR2C showed more robust recruitment of βarr1 than βarr2 (Supplementary Data Figure 6D). As expected, while cluster 1 receptors whose ability to co-localize with β-arrestins at endosomes had not yet been described displayed little signal for endosomal translocation for WT and 3Q β-arrestins, the cluster 2 receptors showed robust signal for recruitment of both WT and 3Q β- arrestins.

Though end-point recruitment of βarr1 to NTSR1 and other cluster 2 GPCRs was largely unaffected by loss of the PIP-binding site, prior NTSR1 experiments had found that loss of PIP binding slowed the kinetics of β-arrestin recruitment (Huang et al., 2020), suggesting PIP_2_ may play a role in the complexes formed with cluster 2 receptors, even when end-point recruitment is unchanged. We fit the rate of β-arrestin translocation to the plasma membrane in response to stimulation for all GPCRs in cluster 2 using our CAAX bystander NanoBiT assay (Figure 1B, top). As was seen for NTSR1, other cluster 2 GPCRs showed a slower association for 3Q than WT (Figure S2A). Though the magnitude of the effect varied across receptors (Figure S2B), these results clearly show that even recruitment to cluster 2 GPCRs is impacted by loss of PIP-binding in β-arrestins.

Together, these results provide several major findings. First, though the tested GPCRs can be divided into two groups, the reliance on PIP-binding for arrestin recruitment is very much a continuum. Generally, GPCRs that co-localize with β-arrestins at endosomes do not require the PIP-binding capacity of β-arrestins for plasma membrane recruitment and are henceforth referred to as PIP-independent GPCRs. Secondly, though PIP-independent GPCRs retained the ability to recruit β-arrestins, the kinetics of recruitment is impaired by loss of PIP binding, suggesting that PIP-mediated interactions likely function to stabilize GPCR-arrestin complexes across all receptors. Finally, while βarr1 and βarr2 behave similarly for most GPCRs, there were exceptions; in much the same way that a continuum of LOF values was observed, this suggests that GPCR-arrestin complexes are incredibly diverse both in their sensitivity to allosteric inputs and possibly their conformational landscape.

### Receptor phosphorylation patterns determine the dependence on PIP-binding by arrestin

The distinction between class A and class B receptors was previously attributed to the presence of suitably positioned clusters of phosphosites in the receptor C-terminus (Oakley et al., 2001). We reasoned that there must be a degree of phosphorylation required to overcome the dependence on arrestin-PIP binding for recruitment to class A receptors. We chose the NTSR1 as a model receptor since WT NTSR1 stably associated with arrestins and the major phosphorylation cluster responsible for this phenomenon was previously established for the rat ortholog (Oakley et al., 2001). Using human NTSR1, we designed a set of phosphorylation-deficient mutants, including both the C-terminus and the third intracellular loop (ICL3) (Figure 2A). In the recent NTSR1-βarr1 structure (Huang et al., 2020), ICL3 was found to be phosphorylated and appeared to make contacts to arrestin, though the role of ICL3 phosphorylation in arrestin recruitment had not been explored. NTSR1 contains four S/T residues in ICL3, three of which are clustered, and 9 S/T residues in its C-terminus, 6 of which are divided into two clusters. To compare the PIP-dependence of NTSR1 phosphorylation mutants (Figure 2A) for their ability to recruit βarr1 to the plasma membrane, we used the CAAX bystander NanoBiT assay (Figure 1A). We first measured cell-surface expression of the NTSR1 constructs and found similar levels (Figure S3A), except for ICL3-4A, which showed slightly reduced expression. Regardless, in this range of receptor expression recruitment signal is saturated with respect to NTSR1 and these minor differences in expression are unlikely to affect the assay response (Figure S3B-C). Though WT NTSR1 is classified as a PIP-independent receptor, NTSR1 phosphorylation mutants could either be classified into cluster 1 or cluster 2 (Figure 2A, Supplementary Data Figure 3), suggesting that particular phosphorylation mutants rendered arrestin recruitment to NTSR1 PIP-dependent. Removal of the two C-terminal phosphorylation site clusters (NTSR1-6A, NTSR1-10A) resulted in a dramatic reduction in arrestin recruitment (Supplementary Data Figure 3), with remaining arrestin recruitment being largely PIP-dependent (Figure 2A). Removal of the ICL3 phosphorylation sites did not affect PIP-dependence (NTSR1-ICL3-4A); neither did removal of the proximal phosphorylation cluster (NTSR1-A^401^VAA), nor removal of any one residue in the distal cluster (NTSR1-TLSA, NTSR1-ALSS, NTSR1-TLAS). However, removal of the distal phosphorylation cluster (NTSR1-A^407^LAA) led to a dramatic reduction in recruitment, and an increase in PIP-dependence, consistent with findings that the distal cluster in the rat ortholog is necessary for stable arrestin association (Oakley et al., 2001). NTSR1-5A, bearing a single C-terminal phosphorylation site in the distal cluster, showed PIP sensitivity comparable to NTSR- ALAA, while NTSR1-4A with two distal cluster phosphorylation sites showing much less PIP- dependence, suggesting that two phosphorylation sites are sufficient to overcome the need for PIP binding. Similarly, NTSR1-TLAA, which differs from NTSR1-5A only in the addition of the proximal cluster of phosphosites exhibits sensitivity between the NTSR1-5A and NTSR1-4A constructs, suggesting that a phosphorylation site from the proximal cluster may offer a partial rescue for the absence of one in the distal cluster.

As the plasma membrane bystander recruitment assay suggested that two phosphorylation sites were necessary to overcome the PIP-dependence on arrestin recruitment, we wondered whether this behavior coincided with the ability of arrestin to be recruited to endosomes. We monitored translocation of arrestin to endosomes using the endosome bystander NanoBiT assay (Figure S1E). As expected, NTSR1-ALAA (Oakley et al., 2001) as well as NTSR1-6A and NTSR1-10A failed to recruit arrestin to endosomes (Figure 2B, Supplementary Data Figure 7). A single C-terminal phosphorylation site (NTSR1-5A) was insufficient to promote arrestin traffic to endosomes; however, two phosphorylation sites in the distal cluster (NTSR1-4A) were sufficient to promote endosomal translocation. There was a further increase in recruitment when the proximal sites were returned (NTSR1-TLSA), suggesting an additional contribution from this region strengthens the interaction between NTSR1 and arrestin. Further support for a contribution from the proximal cluster stems from the difference between NTSR1-5A and NTSR1-TLAA, which differ in the presence of the proximal phosphorylation cluster and show a marked difference in both targeting of arrestin to endosomes, as well as PIP-dependence (Figure 2B). Within the distal cluster, any two phosphorylation sites were sufficient, and having the third present appeared to offer no additional benefit (NTSR1-AVAA compared to NTSR1-ALSS, NTSR1-TLAS and NTSR1- TLSA) (Figure 2B).

Given that two phosphorylation sites in the distal cluster were sufficient for both PIP-insensitivity for plasma membrane recruitment, and recruitment of arrestin to endosomes, we asked whether two phosphorylation sites were also sufficient for receptor internalization. We measured internalization of the NTSR1 constructs in β-arrestin-deficient HEK293 cells where either WT or 3Q βarr1 was reintroduced. WT NTSR1 was robustly internalized by both WT and 3Q βarr1. In contrast, NTSR1-5A showed a significant difference in internalization between WT and 3Q βarr1, while NTSR1-4A showed no difference in internalization between WT and 3Q βarr1. The trend between NTSR1-5A and NTSR1-4A parallels that seen for β2AR and β2AR-V2C (Figure 2C), supporting our finding that two phosphorylation sites are sufficient for robust internalization that is PIP-independent. In addition, the internalization observed for NTSR1-5A by WT βarr1 suggests that the lack of endosome recruitment observed for this construct (Figure 2B) is due to weakened GPCR-βarr interaction and not simply a lack of internalization for this receptor (Figure 2C).

Together, these data show that two suitably positioned phosphorylation sites are sufficient to render β-arrestin recruitment PIP-independent and allow for robust arrestin-dependent internalization as well as support arrestin translocation to endosomes. Furthermore, they show that NTSR1, a receptor that recruits β-arrestin in a PIP-independent manner, can become PIP-dependent by changes in receptor phosphorylation. Given that GPCRs, such as the μOR, have different phosphorylation patterns depending on the stimulating agonist (Just et al., 2013), we speculate that the resulting β-arrestin complexes may have drastically different behavior in cells.

### PIP_2_ binding affects complex stability and tail-core equilibrium in vitro

As PIP-binding was previously suggested to stabilize the interaction between a GPCR and arrestin (Gaidarov et al., 1999), based on experiments in cells, we wanted to explicitly test this *in vitro.* Using NTSR1 as our model receptor, where PIP-binding was not strictly necessary for recruitment in cells, we compared the ability of GRK5 phosphorylated NTSR1 to form a complex with βarr1 (WT or 3Q mutant) in the presence of a soluble PIP_2_ derivative, diC8-PI(4,5)P2 (henceforth PIP_2_), by size-exclusion chromatography (Figure 3A-B) (Huang et al., 2020). While complexing with full-length WT βarr1 led to about 25% complex formation, use of 3Q βarr1 resulted in <5% complex formation (Figure 3C). Use of a C-terminally truncated βarr1 (1-382) led to a more than 2-fold enhancement in complex formation, which was only slightly reduced with the corresponding 3Q arrestin. Using the LOF metric developed to evaluate the impact of PIP-binding on arrestin recruitment in cells, we found that full-length arrestin showed a greater degree of LOF than C-terminally truncated arrestin, suggesting that removal of the arrestin C-terminus is largely able to overcome the impairment in complexing that results from the 3Q mutation (Figure S4A). PIP_2_ affinity *in vitro* was reduced 20x for 3Q βarr1 compared to WT (Figure S4B). Since arrestin activation is understood to proceed via release of its auto-inhibitory C-terminus (Sente et al., 2018; Shukla et al., 2013), we wanted to rule-out the possibility that 3Q βarr1 complexing efficiency is simply reduced due to a lack of arrestin C-terminus release. We designed a Förster Resonance Energy Transfer (FRET) sensor to report on arrestin C-terminus release (Figure S4C): using a cysteine-free βarr1 construct, we introduced two new cysteine residues at positions 12 and 387 – βarr1 (12-387) – to allow for selective labeling of these positions with a suitable dye pair. Given that the expected change in distance between the bound and unbound C-terminus was ∼40 Å (Chen et al., 2017; Kim et al., 2012; Zhuo et al., 2014), we used an AlexaFluor 488/Atto 647N FRET pair, which offers a relatively short Förster radius (R_0_ ∼50 Å). GRK5-phosphorylated NTSR1 robustly displaced the C-terminus of both WT and 3Q βarr1 (12-387) (Figure S4D). Displacement was comparable to that seen for a saturating concentration of a peptide corresponding to the phosphorylated C-terminus of the vasopressin 2 receptor (henceforth V2Rpp) known to completely displace the arrestin C-terminus (Shukla et al., 2013). GRK5- phosphorylated NTSR1 fully displaced the βarr1 C-terminus with 10x greater potency than V2Rpp, suggesting an enhanced affinity for an intact receptor compared to a phosphopeptide alone. These data show that not only does *in vitro* phosphorylated NTSR1 fully displace the arrestin C-terminus, but with higher efficacy than an equimolar concentration of phosphopeptide (even in the presence of unphosphorylated NTSR1), and this is independent of the PIP-binding ability of arrestin.

**Figure 3.**
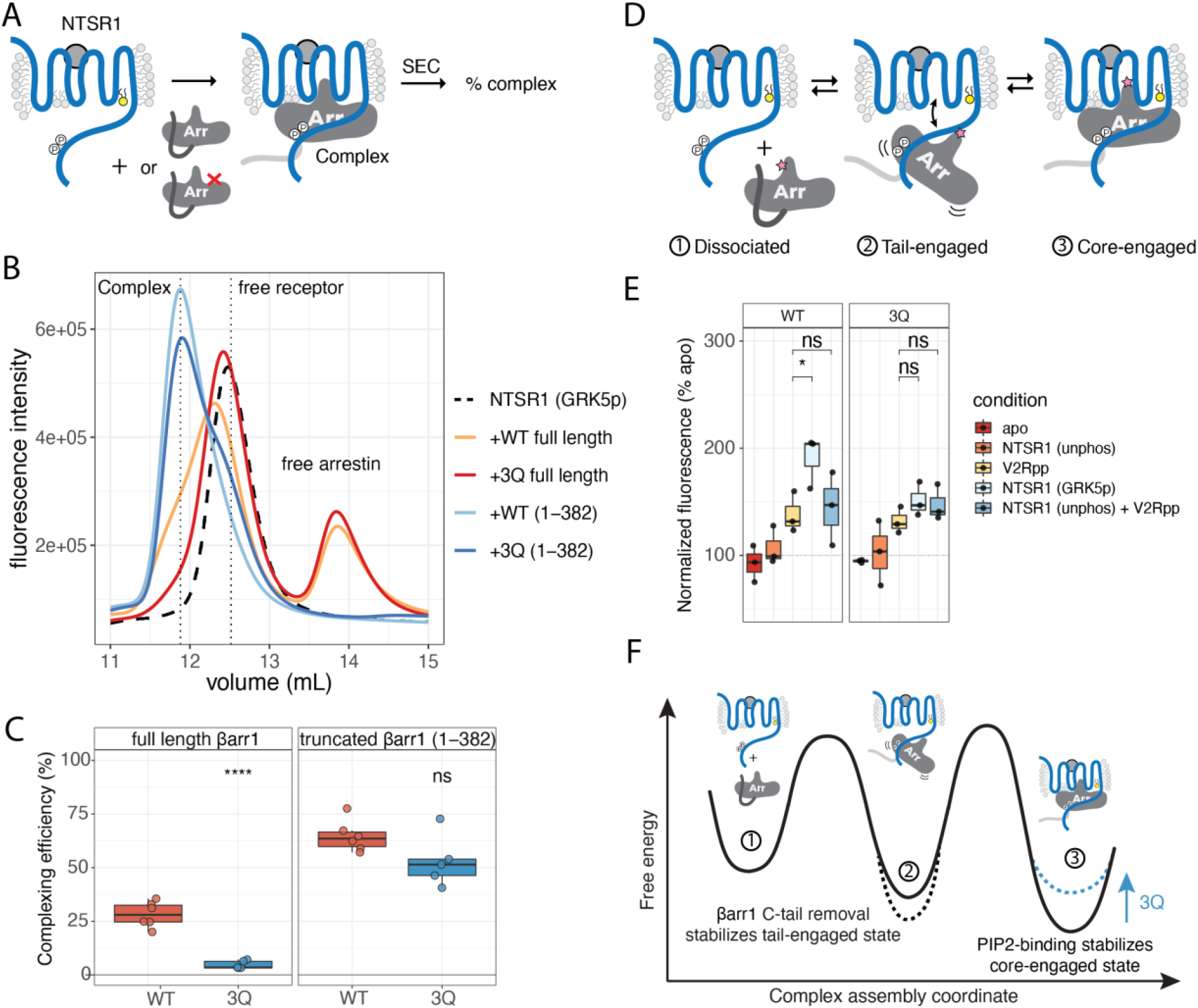
Lipid binding stabilizes core-engaged arrestin complexes. A) cartoon of complexing efficiency assay. Size-exclusion chromatography (SEC) resolves complex from components. B) Representative experiment showing SEC chromatograms with vertical dashed lines indicating free NTSR1, complex, and free arrestins. C) Complexing efficiency, for NTSR1 with indicated arrestins. Boxplots: center line, median; box range, 25–75th percentiles; whiskers denote minimum–maximum values. Individual points are shown (*n*=6 independent experiments). Two- tailed unpaired t-test used to compare conditions. ns: p > 0.05; ****: p ≤ 0.0001. D) Cartoon showing equilibrium of NTSR1-arrestin complex. Pink star denotes L68bim probe used for experiment shown in panel E. E) Bimane spectra for L68bim labeled βarr1 in complex with NTSR1. All NTSR1 samples contained diC8-PI(4,5)P2 (4.1 μM) Boxplots: center line, median; box range, 25–75th percentiles; whiskers denote minimum–maximum values. Individual points are shown (*n*=3 independent experiments). V2Rpp-NTSR1 (GRK5p) and V2Rpp-NTSR1 (unphos) + V2Rpp were compared by two-tailed unpaired t-test. ns: p > 0.05; *: p ≤ 0.05. Apo indicates free arrestin; unphos indicates unphosphorylated receptor and GRK5p indicates GRK5- in vitro phosphorylated receptor. Spectra are normalized to apo (100%) within each experiment and the fluorescence intensity at lambda max was used as the value. F) Free energy diagram illustrating how PIP-binding, by stabilizing the core-engaged state of the NTSR1-arrestin complex slows arrestin dissociation. Loss of the PIP-binding element of arrestin destabilizes the core- engaged state, shifting equilibrium towards the tail-engaged state leading to a higher degree of complex disassembly. Removal of the arrestin C-terminus stabilizes the complex in the tail- engaged state and reduces disassembly even when core-engaged complex is destabilized by lack of PIP-binding.

We reasoned that the reduced complexing efficiency of 3Q βarr1 may be due to differences in the proportion of core-engaged complex being formed (Figure 3D). To test this hypothesis, we used an environmentally sensitive bimane fluorophore (bim) site-specifically installed at L68 (L68bim) on the arrestin finger loop, a region that upon formation of a core-engaged complex with an active GPCR becomes buried within the receptor TM core. Such a sensor had previously been used to report on core-engagement for rhodopsin/arrestin-1 (Sommer et al., 2005, 2006), whereupon receptor core-engagement a blue-shift and an increase in fluorescence emission occurs, owing to the bimane probe moving into a lower polarity environment within the receptor TM core.

While addition of V2Rpp to βarr1 L68bim leads to C-terminus release and a ∼50% increase in bimane fluorescence as seen previously (Latorraca et al., 2020), we speculated that the addition of receptor may further increase this signal (Sommer et al., 2006). We compared the fluorescence changes of βarr1 L68bim (WT or 3Q) upon addition of NTSR1 that was either dephosphorylated or phosphorylated by GRK5 (Figure 3E). In the absence of NTSR1 phosphorylation, there was no increase in fluorescence; however, phosphorylated NTSR1 led to a ∼2-fold enhancement in fluorescence intensity for WT, but a smaller 1.5-fold enhancement for 3Q. The addition of V2Rpp at a saturating concentration to the unphosphorylated NTSR1 did not result in a significant increase over phosphopeptide alone, consistent with the behavior observed for C-terminus release (Figure S4D). Importantly, for the 3Q βarr1, GRK5 phosphorylated NTSR1 did not elicit a response different from V2Rpp alone.

Given that the complex exists as a dynamic equilibrium between three states (Figure 3D): dissociated, tail-bound and core-engaged. We reason that if PIP-binding serves to stabilize the core-engaged state then loss of PIP binding would bias the equilibrium towards a tail-engaged state (Figure 3F), which should have a similar spectroscopic signature to V2Rpp alone. Taken together, these data suggest a model of complex assembly where release of the arrestin C-terminus by the phosphorylated GPCR C-terminus is rapid, and reversible. The resulting tail-bound state is in equilibrium with a core-engaged state, where arrestin-PIP binding stabilized this state and thereby slows dissociation. In the context of full-length arrestin, destabilization of core-engaged state in the 3Q mutant leads to a reduction in complex stability, presumably due to arrestin C-tail-mediated dissociation from the tail-bound state. Consistent with these findings when the arrestin C-terminus is removed the reduced core-engagement of the 3Q mutant does not impact complexing efficiency due to an increased stability of the tail-bound state (as seen in Figure 3C, S4A).

### PIP_2_, in the absence of a GPCR, triggers conformational changes in arrestin

The finding that β-arrestins in CCSs, even in the absence of an associated GPCR, signal through MAPK (Eichel et al., 2016) suggested that arrestin can adopt an active-like conformation without a GPCR C-terminus to displace its own C-terminus. While PIP_2_ was proposed to maintain the membrane association of β-arrestins (Eichel et al., 2018), the impact of this association on the conformational landscape of β-arrestins, and thus their ability to engage downstream signaling partners was unknown (Ranjan et al., 2017). Having shown that PIP-binding affects the dynamics of NTSR1-βarr1 complexes *in vitro*, we wondered whether PIPs in the absence of an associated GPCR could also affect the conformation of βarr1. We compared the effect of PIP_2_ to the V2Rpp for promoting conformational changes in arrestin using FRET and fluorescence reporters on the finger loop, gate loop, and C-terminus (Figure 4A).

**Figure 4.**
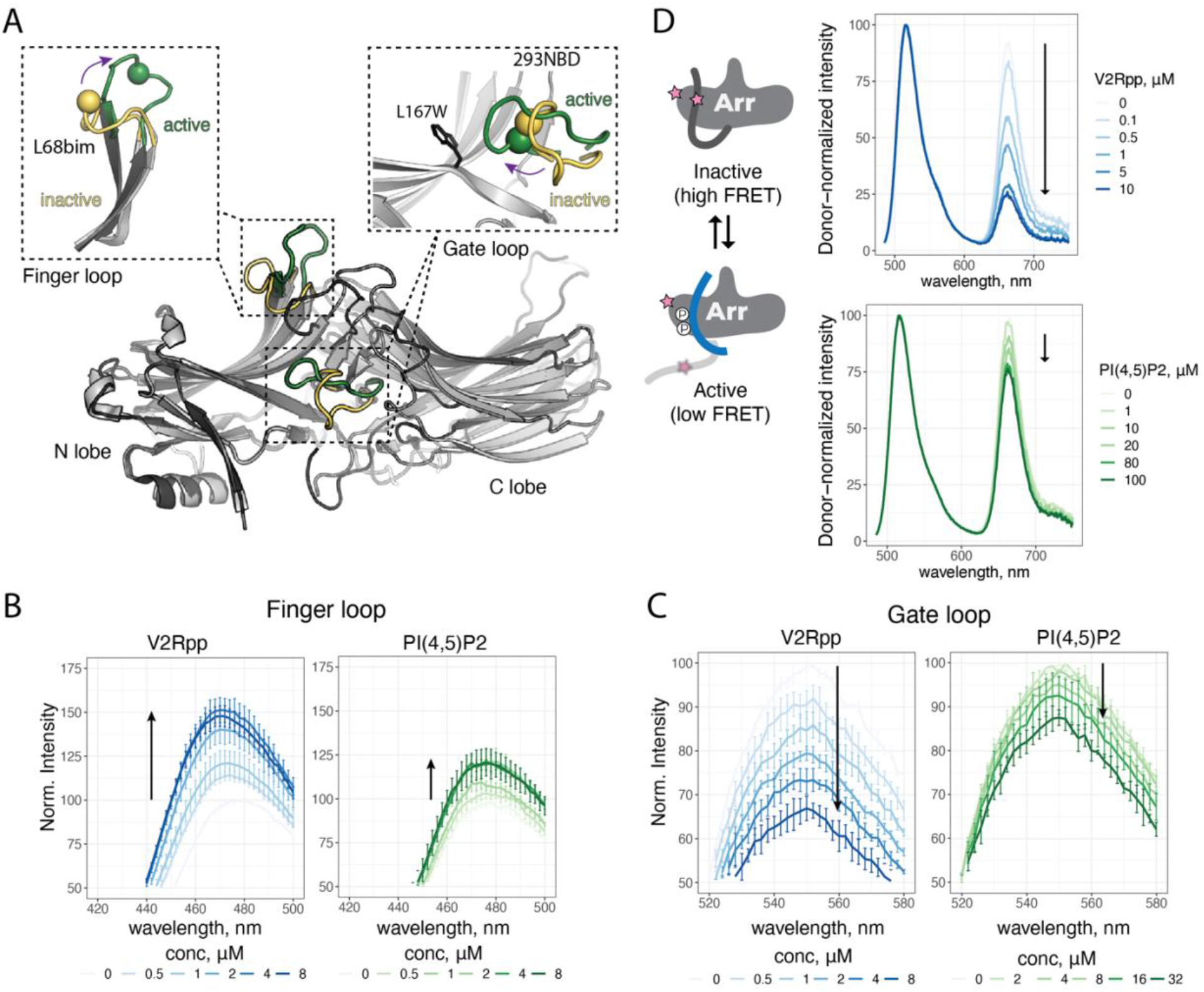
PIP_2_ alone promotes conformational changes in arrestin, including C-terminus movement, but not release. A) overlay of inactive (PDB: 1G4M) [grey] and active (PDB: 4JQI) [black] βarr1. The N and C lobes of βarr1 are indicated. Activation leads to reorganization of several loops, and the gate loop and finger loop are highlighted. Re-orientation of these loops from inactive (yellow) to active (green) can be monitored by site-specific fluorescence spectroscopy. In finger loop inset the sphere denotes Cα L68C which is labeled with bim. In gate loop inset, the sphere denotes Ca L293C which is labeled with NBD. An installed W residue replacing L167 dynamically quenches 293NBD. B) Spectra of bimane labeled (L68C) βarr1 in response to V2Rpp and PIP_2_. Arrow indicates direction of spectral shift with increasing concentration. Values are mean ± SD (*n*=3 independent experiments). Spectra were normalized to the apo condition within a given experiment. C) Spectra of NBD labeled (L167W-L293C) βarr1 in response to V2Rpp and PIP_2_. Arrow indicates direction of spectral shift with increasing concentration. Values are mean ± SD (*n*=3 independent experiments). Spectra were normalized to the apo condition within a given experiment. D) Left, cartoon showing how FRET change is linked to C-terminus release. Right, spectra of AF488/AT647N labeled (A12C-V387C) βarr1 in response to V2Rpp and PIP_2_. Arrow indicates direction of spectral shift with increasing concentration. Spectra were normalized via donor intensity within a given experiment. Data shown are for a representative experiment (*n*=3 independent experiments).

Both the finger loop (Figure 4B, Figure S5A-B) and the gate loop (Figure 4C, Figure S5C-D) showed saturable conformational changes upon addition of PIP_2_ which were smaller than those seen for V2Rpp. Further, the corresponding 3Q mutants did not show PIP_2_-induced conformational changes, though they responded to V2Rpp similarly to WT protein. These data suggest that binding of PIP_2_ to the arrestin C-lobe allosterically promotes conformational changes in key arrestin regions involved in GPCR recognition and activation. As the accepted mechanism for arrestin activation begins with release of its autoinhibitory C-terminus (Sente et al., 2018), we wondered whether these conformational changes were the result of allosterically promoted C-terminus release. Using our βarr1 C-terminus FRET sensor (Figure S4C) we found that PIP_2_ promoted a small movement of the arrestin C-terminus (Figure 4D), but only at concentrations higher than those needed to saturate the responses seen for either the finger or gate loop sensors (Figure 4B-C). As was the case for the other sensors, this FRET change in response to PIP_2_ is absent in the corresponding 3Q mutant (Figure S5E-F). This finding is consistent with recent DEER experiments that found little or no C-terminal displacement for βarr1 with IP6 (Chen et al., 2021). We reason that the conformational changes in the finger and gate loops observed together with the small FRET change in response to PIP_2_ could either be due to a change in the equilibrium of active-inactive βarr1, or a population of an intermediate state of arrestin bearing a change in position or orientation of the arrestin C-terminus within the arrestin N-lobe. As different membrane PIPs serve as markers for different subcellular locations, we measured the ability of other PIPs (Figure S6A) to promote conformational changes in the βarr1 finger loop. Like PI(4,5)P2, PI(3,4)P2, PI(3,5)P2 and PI(3,4,5)P3 all elicited an increase in bimane fluorescence (Figure S6B- F). In contrast, PI(4)P showed a weaker response and PI(3)P and PG did not increase fluorescence of the bimane reporter (Figure S6G-I). Interestingly, these results showed that plasma membrane resident PIPs (Di Paolo and De Camilli, 2006), PI(4,5)P2, PI(3,4)P2, PI(3,4,5)P3 and to a lesser extent PI(4)P were able to promote this conformational change in βarr1, but the early endosomal marker PI(3)P was unable to do so (Figure S6J). PI(3,5)P2 showed a similar effect to other PIP_2_s, but is understood to be rare within cells (Hasegawa et al., 2017). Based on contacts observed in the NTSR1-βarr1 structure (Huang et al., 2020), we speculate that PIPs bearing adjacent phosphates on the inositol ring may be necessary for chelation of K232 and R236/K250 (Figure S6K); however, a phosphate at the 4-position is sufficient to coordinate K250 and R236, explaining the small effect seen for PI(4)P. Together, these data show that different PIP_2_ derivatives are capable of promoting conformation changes in βarr1, while PIPs bearing a single phosphate do not, raising the possibility of compartment-specific differences in the behavior of GPCR-arrestin complexes.

### PIP_2_ increases the population of active arrestin

While our fluorescence experiments support PIP_2_-promoted conformational changes consistent with arrestin activation, the lack of C-terminus release raised questions of whether these conformational changes truly reflected an increase in the population of active arrestin, as would be detected by arrestin binding partners. While the active form of arrestin is understood to mediate signaling via interactions with a number of protein partners, including MAPK, ERK, SRC (Ranjan et al., 2017; Reiter et al., 2012), there has been speculation that the binding of a particular partner might be mediated by a distinct arrestin conformation. We reasoned that the global activation state of arrestin could be probed using an engineered Fab (Fab30), which has a high-affinity for the active (V2Rpp-bound) state of βarr1 (Shukla et al., 2013). Fab30 has found utility in a number of structural studies (Lee et al., 2020; Nguyen et al., 2019; Shukla et al., 2013; Shukla et al., 2014; Staus et al., 2020), functional studies (Cahill et al., 2017; Ghosh et al., 2019; Kumari et al., 2016; Latorraca et al., 2020; Thomsen et al., 2016) and more recently it has been adapted as a single-chain intrabody (IB30) for the detection of active βarr1 in cells (Baidya et al., 2020a; Baidya et al., 2020b).

We used Surface Plasmon Resonance (SPR) to measure binding of Fab30 to immobilized βarr1 (Figure 5A). To confirm the immobilized arrestins behave as expected, we tested binding of V2Rpp and Fab30+V2Rpp (Figure 5B, Figure S8, Supplementary Data Tables 2-3). Though selected for binding to the V2Rpp-bound state of βarr1, Fab30 bound to βarr1 weakly in the absence of V2Rpp, presumably due to a small equilibrium population of active-like arrestin (Latorraca et al., 2018). Interestingly, binding was enhanced when Fab30 was co-injected with PIP_2_ (Figure 5B). This suggested that PIP_2_ increased the proportion of arrestin in an active-like state which can be recognized by Fab30, consistent with our fluorescence experiments that support PIP_2_ playing a role in arrestin activation. We compared the effect of different additives on Fab30 binding to WT βarr1, but also a βarr1 3Q mutant, and the pre-activated C-terminally truncated βarr1(1-382) (Kim et al., 2013). At 1 μM, Fab30 showed 10.2 ± 0.9% (of maximal) binding to WT βarr1, compared to 56.8 ± 2.0% binding for βarr1 (1-382) (Figure 5C). This suggests that Fab30 binding is favored by a conformation accessible to WT βarr1, but greatly enhanced by removal of the arrestin C-terminus. When Fab30 is co-injected with a saturating concentration of PIP_2_ (40 μM), binding to WT βarr1 increased more than 3-fold, to 33.9 ± 1.8%, compared to Fab30 alone. PIP_2_ had a smaller effect on the pre-activated βarr1(1-382), but still increased binding from 56.8% to 65.9 ± 0.8%. Titration experiments showed that specific enhancement of Fab30 binding in the presence of PIP_2_ was most pronounced for WT βarr1 (Figure S7G-L). While all three arrestin constructs showed an increase in Fab30 binding in the presence of PIP_2_, the degree of binding enhancement drastically shifted for WT βarr1 above the K_d_ for Fab30, but not for either 3Q or (1-382) βarr1 (Supplementary Data Figure 8). This suggests that while PIP_2_ enhanced the population of active-like βarr1, Fab30 binding remains rate-limiting. Since removal of the βarr1 C-terminus abrogates the PIP_2_-enhancement of Fab30 binding, we reason that PIP_2_ acts in *cis* with C-terminal displacement, consistent with our FRET experiments that showed a PIP_2_-induced movement of the βarr1 C-terminus. To determine whether this effect was specific for PIP_2_, we compared the ability of PG and PI(3)P to enhance binding of Fab30 . Both showed a small enhancement in Fab30 binding, but significantly less than that seen with PIP_2_ (Figure 5C). Further, βarr1 3Q showed no difference between PG, PI(3)P and PIP_2_, suggesting that while anionic lipids weakly increase Fab30 binding to βarr1, PIP_2_ was unique in affecting a specific increase in Fab30 binding. Both PG and PI(3)P did not enhance Fab30 binding to βarr1 (1-382).

**Figure 5.**
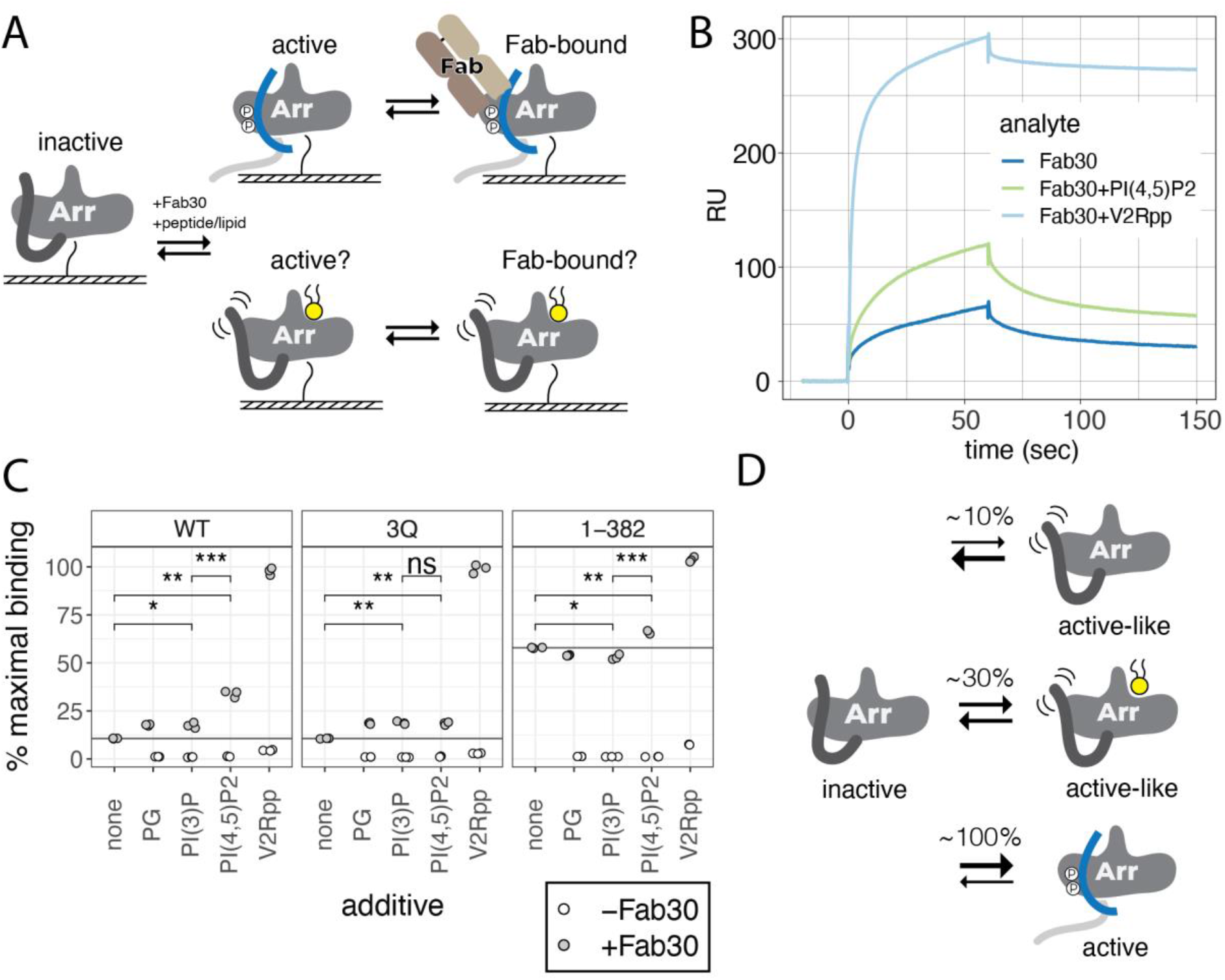
PIP_2_ enhances Fab30 binding to βarr1. A) Cartoon of surface plasmon resonance (SPR) experiments, where βarr1 is immobilized via N-terminal biotinylation and a Fab30 binder is injected in the presence of absence of PIP_2_ or V2Rpp. B) Representative sensogram for SPR binding experiment. With WT βarr1 immobilized, Fab30 (1 μM) was injected alone or together with V2Rpp (40 μM) or diC8-PI(4,5)P2 (40 μM). The shown sensogram is representative of the outcome seen for independent experiments (*n*=3). Dissociation/regeneration phase not shown. C) Binding of Fab30 to immobilized arrestin constructs in the presence of different additives. Maximum binding is defined based on normalization of the observed response to the amount of arrestin immobilized for each construct. Additives: diC8-PG (40 μM), diC8-PI(3)P (40 μM), diC8- PI(4,5)P2 (40 μM) and V2Rpp (40 μM) were mixed with Fab30 (1 μM) and injected together. Points reflect independent measurements; open points represent the binding observed for the additive in the absence of Fab30. Fab30 binding was compared using a two-tailed unpaired t-test. ns: p > 0.05; *: p ≤ 0.05; **: p ≤ 0.01; ***: p ≤ 0.001. D) The proportion of active-like βarr1 increases in the presence of PIP_2_.

Based on these data we propose that spontaneous activation of arrestin to an active-like state capable of binding Fab30 is possible but rare in the absence of arrestin inputs (Figure 5D). V2Rpp dramatically shifts the equilibrium towards the active-state by displacement of the arrestin C-terminus, and removal of the arrestin C-terminus alone is sufficient to greatly enhance the active- population, even in the absence of V2Rpp or PIP_2_. Unlike V2Rpp, which displaces the arrestin C-terminus, PIP_2_ is unable to displace the arrestin C-terminus directly, but able to allosterically move it. While PIP_2_ may stabilize the same active state of arrestin achieved with V2Rpp, albeit to a lesser extent, it may also act to stabilize an active-like state of arrestin that is on-pathway towards activation and capable of binding Fab30, though to a lesser extent than V2Rpp-bound βarr1. Further studies will be necessary to distinguish these possibilities.

## Conclusions

Our results reveal new molecular details underpinning the regulation of arrestin recruitment to GPCRs, and how spatial and temporal control of GPCR-β-arrestin complexes may occur within a cell.

Our findings offer a molecular basis for understanding the phenotypic classification of GPCRs into “class A” or “class B” for arrestin recruitment. In our model (Figure 6), we refer to “class A” and “class B” GPCRs as “PIP-dependent” and “PIP-independent”, respectively. “PIP-dependent” GPCRs (Figure 6, left) require the coincident detection of membrane PIPs for recruitment to an activated and phosphorylated GPCR. We speculate that this is due to an insufficiency in phosphorylation of these receptors, requiring either an allosteric priming of C-terminus release by plasma membrane PIPs, or the simultaneous action of both phosphate-mediated contacts and PIP-mediated contacts to form a sufficiently long-lived complex for effective receptor desensitization, sequestration and internalization. As some PIP-dependent GPCRs can recruit arrestin in a C-terminus-independent manner, we consider that release of the arrestin C-terminus may not be necessary for arrestin function in the context of these receptors. A further trait of these PIP-dependent GPCRs is that they exhibit, to a varying degree, the “catalytic activation” phenotype (Eichel et al., 2018) wherein arrestin, after recruitment to an active GPCR, loses association with the GPCR but remains at the plasma membrane and concentrates at CCSs. This can be explained by the increasing concentration gradient of PIP_2_ leading into the CCS (Sun et al., 2007) along with our biophysical evidence that PIP_2_ promotes conformational transitions associated with activation. Once a GPCR cargo has been translocated into a CCS, clathrin-mediated endocytosis (CME) proceeds and PIP_2_ levels drop. We suggest that this may serve as the timing component for arrestin dissociation from these PIP-dependent GPCRs (Zhang et al., 1999). Presumably, once arrestin has dissociated, the receptor is susceptible to dephosphorylation, and upon arrival at early endosomes is able to be sorted for rapid recycling (Krueger et al., 1997). In contrast, “PIP-independent receptors” (Figure 6, right panel) possess phosphorylation sites which alone can promote a stable association with arrestin, without the need for membrane PIPs. Since PIP-binding is not necessary to maintain the GPCR-arrestin association, arrestin co-localizes with these receptors at endosomes. Whether this co-localization is the product of PIP-independent GPCRs being able to recruit β-arrestins when at endosomes or by forming a sufficiently stable complex to allow for co-trafficking from the plasma membrane without exchange remains to be shown.

**Figure 6.**
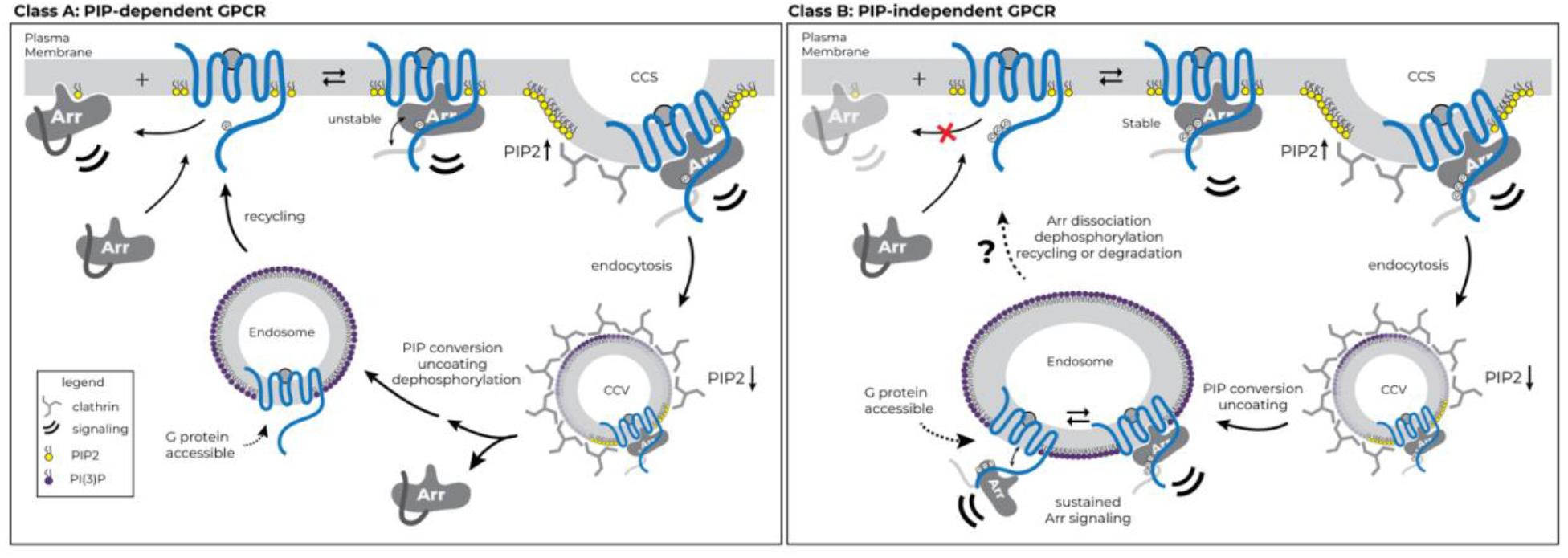
Model for phosphoinositide regulation of GPCR-β-arrestin complex assembly and disassembly. GPCRs stratify into two groups with respect to the strength of their interaction with β-arrestins: one group requires an interaction between β-arrestin and PIP_2_ at the plasma membrane for recruitment (PIP-dependent), while the other does not (PIP-independent). In the case of PIP-dependent GPCRs, arrestin engagement is unstable and can result in dissociation of arrestin from the receptor, while maintaining an association with the plasma membrane (left panel). PIP_2_ is enriched at CCSs and in both cases complex assembly can occur. During endocytosis, PIP_2_ is depleted and for PIP-dependent GPCRs, the loss of this PIP_2_ contact may facilitate dissociation of arrestin thereby allowing for receptor recycling. In contrast, a PIP- independent GPCR will retain the interaction with arrestin even once PIP_2_ is depleted owing to the strong phosphorylation-dependent interactions; however, the full-engaged state of the complex is less stable in endosomes than at the plasma membrane, thereby allowing further G protein engagement to occur.

One question this model raises is: if PIP_2_ promotes partial activation of β-arrestins, why is it that β-arrestins are not basally associated with the plasma membrane? Consistent with our SPR experiments, and the finding that GPCR C-terminal phosphorylation is required for arrestin accumulation at CCSs (Eichel et al., 2018), we speculate that PIP_2_ binding may occur when arrestin transitions to an active-like conformation and stabilize this state. We speculate that in cells, engagement with a GPCR may be necessary to facilitate PIP_2_ binding. GPCRs have been shown to associate with PIP_2_ in the local membrane environment (Song et al., 2019; Yen et al., 2018), and in doing so may act to “load” PIP_2_ onto the arrestin, which either can remain associated with the GPCR or diffuse along the membrane.

These data suggest that while PIP-mediated contacts are not necessary to maintain association, they likely affect the equilibrium of core vs. tail-engaged states of the complex. Tail-engagement has been shown to be sufficient for MAPK signaling downstream of β-arrestin (Kumari et al., 2017). We speculate that this shift in equilibrium, particularly in the context of endosomes defined by PI(3)P, may explain how PIP-independent receptors, such as V2R and PTH1R are able to engage and signal through both β-arrestin and G proteins simultaneously in a so-called “megaplex” assembly (Nguyen et al., 2019; Thomsen et al., 2016).

To-date four structures of GPCR-βarr1 complexes have been described, all of which show arrestin in a core-engaged state (Huang et al., 2020; Lee et al., 2020; Staus et al., 2020; Yin et al., 2019), but only one had PIP_2_ bound at the interface (Huang et al., 2020). Interestingly, this NTSR1-βarr1 complex with PIP_2_ bound used the native NTSR1 C-terminus and did not use Fab30 to stabilize the complex. We speculate that Fab30 plays a particularly important role in stabilizing the receptor core-engaged complex (Shukla et al., 2014).

Overall, our data offer a parsimonious explanation for several phenotypic behaviors observed for GPCR-β-arrestin complexes and provide a biophysical framework for understanding the interplay between phosphorylation-mediated and PIP-mediated contacts in complex assembly. A reliance on PIPs for arrestin recruitment offers a robust solution for recruitment of arrestin to receptors with spatial control, and temporal precision. Given the interplay between PIP-dependent recruitment and phosphorylation, we believe that distinct signaling outcomes may not only be due to differences in phosphorylation alone (Latorraca et al., 2020), but rather that these differences may be further fine-tuned by membrane PIPs that are present in distinct subcellular locations, adding yet another layer of complexity to our understanding of GPCR signaling.

## Acknowledgements

We thank Betsy White for technical laboratory assistance; Shoji Maeda for providing purified Fab30 protein; Weijiao Huang for assistance with NTSR1 expression and purification; Daniel Hilger for purified BirA enzyme; Yoon Seok Kim and Eamon Byrne for assistance with FSEC experiments, Kayo Sato, Yuko Sugamura, Shigeko Nakano and Ayumi Inoue for plasmid construction and cell based GPCR assays. We thank R. Scott Prosser for additional comments on the manuscript. This work was supported in part by National Institutes of Health grants R01NS028471 (B.K.K.), R01 AI125320 (K.C.G), R01DA010711 and R01DA012864 (M.vZ.). Additional support to both K.C.G and B.K.K. is provided by the Mathers Foundation. B.K.K. is a Chan-Zuckerberg Biohub Investigator. J.J. is a Damon Runyon Fellow supported by the Damon Runyon Cancer Research Foundation (DRG-2318-18). B.B.R. is a recipient of an American Heart Association Predoctoral Fellowship (19PRE34380570). F.M. H. is a recipient of a Marie Skłodowska-Curie Individual Fellowship from the European Union’s Horizon 2020 research and innovation programme (grant agreement No. 844622) and an American Heart Association postdoctoral fellowship (19POST34380839). M.M. was supported by an American Heart Association postdoctoral fellowship (17POST33410958). A.I. was funded by the PRIME 19gm5910013, the LEAP 20gm0010004 and the BINDS JP20am0101095 from the Japan Agency for Medical Research and Development (AMED); KAKENHI 17K08264, 21H04791 and 21H051130 from by the Japan Society for the Promotion of Science (JSPS); JST Moonshot Research and Development Program JPMJMS2023 from Japan Science and Technology Agency (JST); Daiichi Sankyo Foundation of Life Science; Takeda Science Foundation; Ono Medical Research Foundation; The Uehara Memorial Foundation.

## Author Contributions

Conceptualization, J.J. and A.I.; Methodology, J.J., B.B-R., A.I.; Software, J.J., F.M.H.; Formal Analysis, J.J., F.M.H.; Investigation, J.J., R.K., B.B-R., D.H.S., M.M., A.I.; Resources, M.M., K.K., Data Curation, J.J., F.M.H., A.I.; Writing – Original Draft, J.J.; Writing – Review & Editing, J.J., R.K., B.B-R., D.H.S., F.M.H., K.K., M.vZ., K.C.G., A.I., B.K.K.; Visualization, J.J., F.M.; Supervision, K.C.G., M.vZ., A.I., B.K.K.; Funding Acquisition, K.C.G., M.vZ., A.I., B.K.K.

## Methods

### Plasmids

For cell-based assays, we used human, full-length GPCR plasmids cloned into the pCAGGS vector or the pcDNA3.1 vector derived from a previous study (Inoue et al., 2019). GPCR constructs were N-terminally FLAG epitope-tagged when they were intended to compare with cell surface expression levels. Specifically, NTSR1 was fused to the N-terminal FLAG epitope tag with a linker (M<u>DYKDDDDK</u>GTELGS; the FLAG epitope tag is underlined) and inserted into the pcDNA3.1 vector. β2AR and μOR were fused to the N-terminal FLAG epitope tag with a preceding HA-derived signal sequence and a flexible linker (MKTIIALSYIFCLVFA<u>DYKDDDDK</u>GGSGGGGSGGSSSGGG) and inserted into the pCAGGS vector. Unless otherwise noted, other GPCR constructs were untagged. For the bystander NanoBiT-based β-arrestin assays, human full-length β-arrestin (β-arrestin1 or 2; WT or 3Q) was N-terminally SmBiT-fused with the flexible linker (M<u>VTGYRLFEEIL</u>GGSGGGGSGGSSSGG; the SmBiT is underlined) and inserted into the pCAGGS vector (SmBiT-β-arrestin) (Baidya et al., 2020a). For the plasma membrane-localizing tag, LgBiT was C-terminally fused to the CAAX motif derived from human KRAS (SSSGGGKKKKKKSKTKCVIM) through the same flexible linker (LgBiT-CAAX). For the endosome-localizing tag, LgBiT was N-terminally fused with the human Endofin FYVE domain (amino-acid regions Gln739-Lys806) again through the same flexible linker (Endo-LgBiT). For the direct NanoBiT-based β-arrestin assay, human full-length β-arrestin was N-terminally LgBiT-fused with the same flexible linker and inserted into the pCAGGS vector (LgBiT-β-arrestin). GPCRs were C-terminally SmBiT-fused with the flexible linker (GGSGGGGSGGSSSGG<u>VTGYRLFEEIL</u>; the SmBiT is underlined) and inserted into the pCAGGS vector (GPCR-SmBiT).

### Peptides

The V2Rpp peptide (ARGRpTPPpSLGPQDEpSCpTpTApSpSpSLAKDTSS) was obtained by custom peptide synthesis (Tufts University Core Facility). Fab30 was expressed and purified as previously described (Shukla et al., 2013). The concentration of V2Rpp stocks were determined by reaction with Ellman’s reagent as previously described (Latorraca et al., 2020).

### NanoBiT-β-arrestin recruitment assays

β-arrestin recruitment to the plasma membrane was measured by the bystander NanoBiT-β- arrestin assays using the SmBiT-β-arrestin and the LgBiT-CAAX constructs. HEK293A cells (Thermo Fisher Scientific) were seeded in a 6-cm culture dish (Greiner Bio-One) at a concentration of 2 x 10^5^ cells per ml (4 ml per dish hereafter) in DMEM (Nissui Pharmaceutical) supplemented with 10% FBS (Gibco), glutamine, penicillin, and streptomycin, one day before transfection. The transfection solution was prepared by combining 5 μl of polyethylenimine solution (1 mg/ml) and a plasmid mixture consisting of 100 ng SmBiT-β-arrestin, 500 ng LgBiT-CAAX and 200 ng of a test GPCR construct in 200 µl of Opti-MEM (Thermo Fisher Scientific). For the NTSR1 titration experiment, diluted volume of the FLAG-NTSR1 plasmid (13 ng to 200 ng) was transfected with 100 ng SmBiT-β-arrestin and 500 ng LgBiT-CAAX with a balanced volume of the pcDNA3.1 vector (total plasmid volume of 800 ng). After an incubation for one day, the transfected cells were harvested with 0.5 mM EDTA-containing Dulbecco’s PBS, centrifuged, and suspended in 2 ml of Hank’s balanced saline solution (HBSS) containing 0.01% bovine serum albumin (BSA fatty acid–free grade, SERVA) and 5 mM HEPES (pH 7.4) (assay buffer). The cell suspension was dispensed in a white 96-well plate (Greiner Bio-One) at a volume of 80 μl per well and loaded with 20 µl of 50 µM coelenterazine (Carbosynth), diluted in the assay buffer. After 2 h incubation at room temperature, the plate was measured for its baseline luminescence (SpectraMax L, 2PMT model, Molecular Devices). Thereafter, 20 μl of 6x ligand serially diluted in the assay buffer were manually added. The ligand used was dependent on the GPCR expressed, as described in Supplementary Data Table 1. The plate was immediately read for the second measurement as a kinetics mode and luminescence counts recorded for 15 min with an accumulation time of 0.18 sec per read and an interval of 20 sec per round. β-arrestin endosomal translocation was measured by following the same procedure as described above but using the SmBiT-β-arrestin and the Endo-LgBiT constructs. Similarly, direct recruitment was measured by the same protocol as described above but using LgBiT-β-arrestin (500 ng) and C-terminally fused- SmBiT GPCR (500 ng) constructs. For every well, the recorded kinetics data were first normalized to the baseline luminescence counts.

### Analysis of cell-based recruitment data

NanoBiT data were analyzed by converting kinetic data into concentration-response data by determining an average fold-change (relative to signal pre-stimulation) from 10-15 minutes post-agonist addition. At least three independent experiments were performed for each receptor- sensor combination. Concentration-dependent data from two technical replicates for each independent experiment were collectively fit to a four-parameter log logistic function (LL2.4) provided in the drc package (v 3.0-1) of the statistical environment R. This equation, of the form: 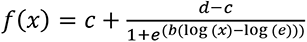 provides pre- and post-transition values, c and d, respectively, that define the amplitude response for that assay. Cutoffs for bystander NanoBiT experiments were determined as based on a limit of detection of 3s over the response of mock-transfected cells. Amplitude values were defined as amplitude = top – bottom of fit, and amplitude error was calculated as 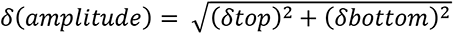. Converting amplitude to LOF for each assay was based on the formula:1 − *amplitude*(*3Q*)/*amplitude*(*WT*). Errors for LOF were calculated as: . In cases where a fit failed to 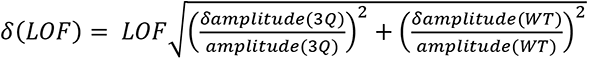 converge due to weak recruitment, these amplitudes and errors were set to zero. Recruitment of βarr1 (3Q) to D1R in both plasma membrane bystander (CAAX) and direct recruitment which was set to zero. The error amplitude for βarr1 (3Q) endosome translocation assay with D1R was also set to zero. The error amplitude for βarr1 (3Q) endosome translocation assay with S1PR1 was set to zero, and the “top” value of the fit was set to 1.2 based on manual inspection. K-means clustering was performed using pre-built functions in the tidyverse package (v 1.3.1) of R. The number of clusters was varied from 1 to 10 and an elbow plot of within cluster sum of squares vs k suggested 2 clusters fit the data well.

For recruitment kinetics, luminescence fold-change was plotted against time, and the values from zero to five minutes (initial rate) were fit to a logistic function of the form: 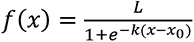, where L is the curve’s maximum value, x_0_ is the value of the sigmoid midpoint and k is the logistic growth rate. Fitting was done using the self-starting SSlogis four parameter nls function in the tidyverse package (v 1.3.1) of R.

### GPCR internalization assay

GPCR internalization assays was performed as described previously with minor modifications (Grundmann et al., 2018). Δβarr1/2 double knockout (DKO) cells, previously described (O’Hayre et al., 2017), were seeded in 6-cm dishes at concentration of 2 x 10^5^ cells/ml (4 mL per dish) and cultured for 1 day before transfection. The cells were transfected with 1 μg of the N-terminally FLAG-tagged NTSR1 or the β2AR construct, along with 200 ng of the WT or 3Q βarr1 or empty plasmid, using PEI transfection reagent as described above. After 1-day culture, the transfected cells were harvested by EDTA-PBS and HEPES-HBSS and, following centrifugation, the cells were suspended in 500 μL of 0.01% BSA-containing HEPES-HBSS. The cell suspension was dispensed in a 96-well V-bottom plate (100 µL per well) and mixed with 100 μL of 2× GPCR solution ligand (2 μM neurotensin for FLAG-NTSR1 or 20 μM Isoproterenol (Sigma-Aldrich) for FLAG-β2AR). After 30-min incubation in a CO_2_ incubator, the plate was centrifuged at 1,500 g for 5 min and the cells were washed twice with D-PBS. The cell pellets were suspended in 2% goat serum- and 2 mM EDTA-containing D-PBS (blocking buffer; 100 µL per well) and incubated for 30 min on ice. After centrifugation, the cells were stained with anti-FLAG-epitope tag monoclonal antibody (Clone 1E6, FujiFilm Wako Pure Chemicals; 10 µg mL-1 in the blocking buffer; 25 μL per well) for 30 min on ice. After washing with D-PBS, the cells were labeled with a goat anti-mouse IgG secondary antibody conjugated with Alexa Fluor 647 (Thermo Fisher Scientific; 10 µg mL^-1^ dilution in the blocking buffer; 25 μL per well) for 15 min on ice. The cells were washed once with D-PBS, resuspended in 100 μL of 2 mM EDTA-containing-D-PBS and filtered through a 40 μm filter. The fluorescently labeled cells (approximately 20,000 cells per sample) were analyzed by the EC800 flow cytometer (Sony). Fluorescent signal derived from Alexa Fluor 647 was recorded in the FL3 channel. Mean fluorescence intensity (MFI) from all of the recorded events was analyzed by a FlowJo software (FlowJo) and used for statistical analysis.

### Cell-surface expression analysis by flow cytometry

HEK293A cells were seeded in a 6-well culture plate at concentration of 2 x 10^5^ cells/ml (2 mL per dish) and cultured for 1 day before transfection. The cells were transfected with 1 μg of N-terminally FLAG-tagged GPCR construct using PEI transfection reagent as described above and cultured for 1 day. The cells were collected by adding 200 μl of 0.53 mM EDTA-containing Dulbecco’s PBS (D-PBS), followed by 200 μl of 5 mM HEPES (pH 7.4)-containing Hank’s Balanced Salt Solution (HBSS). The cell suspension was transferred to a 96-well V-bottom plate in duplicate and fluorescently labeled with the anti-FLAG epitope tag antibody and a goat anti-mouse IgG secondary antibody conjugated with Alexa Fluor 488 (Thermo Fisher Scientific, 10 μg per ml diluted in the blocking buffer) as described above. Live cells were gated with a forward scatter (FS-Peak-Lin) cutoff at the 390 setting, with a gain value of 1.7 and fluorescent signal derived from Alexa Fluor 488 was recorded in the FL1 channel. For each experiment, the MFI value of mutants was normalized to that of WT performed in parallel.

### cAMP desensitization

HEK293 Δβarr1/2 (DKO) cells that endogenously express β2AR were seeded into 6-well plates and transiently transfected after 24 hours with mApple, βarr2-mApple, or βarr2(3Q)-mApple. Twenty-four hours after transfection, cells were transduced with CMV cADDis Green Upward cAMP sensor according to manufacturer instructions without addition of sodium butyrate (Montana Molecular #U0200G) and seeded in triplicate in a black clear-bottom 96-well plate (Corning cat# 3340). Twenty-four hours after transduction, the cells were washed once with 37°C assay buffer [135 mM NaCl, 5 mM KCl, 0.4 mM Mg2Cl, 1.8 mM CaCl2, 5 mM glucose, 20 mM HEPES pH 7.4], loaded into the pre-warmed 37°C plate reader (Biotek Synergy H4), and equilibrated for five minutes. Prior to beginning the kinetic assay, mApple was read using monochromoters set to Ex:568/9.0 and Em:592/13.5. Then cADDis was read using monochromoters set to Ex:500/9.0 and Em:530/20.0. Three cADDis timepoints were collected to establish baseline, the plate was ejected, isoproterenol in 37°C assay buffer was added to a final concentration of 100 nM, and the plate was returned to continue collection. Thirty minutes after isoproterenol addition, 3-isobutyl-1-methylxanthine (IBMX) and forskolin (Fsk) in 37°C assay buffer were added to a final concentrations of 300 μM and 10 μM respectively. Responses were averaged across technical replicates, normalized to the maximum Fsk/IBMX response, and then averaged across independent experiments. Expression levels for cADDis and βarr2 were normalized based on fluorescence.

### Western blotting

HEK293A cells were transfected with the SmBiT-β-arrestin and the LgBiT-CAAX constructs by following the procedure described in the NanoBiT-based β-arrestin assay. After 1-day culture, the transfected cells were lysed by SDS-PAGE sample buffer (62.5 mM Tris-HCl (pH 6.8), 50 mM dithiothreitol, 2% SDS, 10% glycerol and 4 M urea) containing 1 mM EDTA and 1 mM phenylmethylsulfonyl fluoride. Lysates derived from an equal number of cells were separated by 8% SDS-polyacrylamide gel electrophoresis. Subsequently, the proteins were transferred to PVDF membrane. The blotted membrane was blocked with 5% skim milk-containing blotting buffer (10 mM Tris-HCl (pH 7.4), 190 mM NaCl and 0.05% Tween 20), immunoblot with primary (1 µg per mL, unless otherwise indicated) and secondary antibodies conjugated with horseradish peroxidase (1:2000 dilution). Primary antibodies used in this study were: anti-β-arrestin1 (rabbit monoclonal; CST, #12697, D8O3J), anti-β-arrestin2 antibody (rabbit monoclonal; CST, #3857, C16D9) and anti-α-tubulin antibody (mouse monoclonal, clone DM1A; Santa Cruz Biotechnologies, sc-32293; 1:2000 dilution). Secondary antibodies were anti-rabbit IgG (GE Healthcare, NA9340) and anti-mouse IgG (GE Healthcare, NA9310). Membrane was soaked with an ImmunoStar Zeta reagent (FujiFilm Wako Pure Chemical). Chemiluminescence image of the membrane was acquired, and band intensity was analyzed with Amersham Imager 680 (Cytiva).

### NTSR1 expression and purification

Full length human NTSR1 was modified with an N-terminal Flag tag followed by an octa-histidine tag and cloned into pFastBac1 vector. NTSR1 was expressed in Sf9 insect cells (Expression Systems) using a FastBac-derived baculovirus. Cells were infected at a density of 4x10^6^ cells/mL and harvested 60 hrs post infection. Cells were lysed in hypotonic buffer (10 mM HEPES, pH 7.4, and protease inhibitors) and solubilized at 4 °C for 2 hours in a buffer containing 1% lauryl maltose neopentyl glycol (LMNG, Anatrace), 0.1% cholesteryl hemisuccinate tris salt (CHS, Steraloids), 0.3% sodium cholate (Sigma), 20 mM HEPES 7.4, 500 mM NaCl, 25% glycerol, iodoacetamide (to cap cysteine residues) and protease inhibitors. Insoluble debris was removed by centrifugation and the supernatant was incubated with Ni-NTA (Qiagen) resin for 1 hour at 4 °C. The resin was washed in batch with buffer containing 0.01% LMNG, 0.001% CHS, 0.003% sodium cholate, 20 mM HEPES pH 7.4, 500 mM NaCl, 10 mM imidazole and eluted with the same buffer supplemented with 200 mM imidazole, 2 mM CaCl_2_ and 10 μM NTS_8-13_ (Acetate salt, Sigma). The eluate was loaded onto M1 FLAG immunoaffinity resin and washed with buffer containing 0.01% LMNG, 0.001% CHS, 0.003% sodium cholate, 20 mM HEPES pH 7.4, 500 mM NaCl, 10 mM imidazole, 0.1 μM NTS_8-13_ and 2 mM CaCl_2_. The receptor was eluted with buffer containing 100 mM NaCl, 20 mM HEPES pH 7.4, 0.005% LMNG, 0.005% CHS, 1 μM NTS_8-13_, 0.2 mg/mL flag peptide (DYKDDDDK) and 5 mM EDTA. Elution fractions containing receptor were pooled and subjected to polishing by SEC on a Superdex 200 Increase 10/300 GL column (GE Healthcare) in 20 mM HEPES, pH 7.4, 100 mM NaCl, 0.0025% LMNG, 0.00025% CHS, and 0.1 μM NTS_8-13_. Peak fractions were pooled and concentrated to 200 μM and aliquots were flash-frozen and stored at -80 °C until use.

### GRK5 expression and purification

Full length human GRK5 was modified with a C-terminal hexa-histidine tag and cloned into pVL1392 vector for baculovirus production. GRK5 was expressed and purified as previously published (Beyett et al., 2019). Briefly, Sf9 insect cells (Expression Systems) were infected with a BestBac-derived baculovirus at a density of 3.5 x 10^6^ cells/mL and harvested 48 hours post infection. Cells were resuspended, lysed by sonication and the supernatant was applied to Ni-NTA resin. The resin was washed with lysis buffer and GRK5 eluted with lysis buffer supplemented with 200 mM imidazole. The combined eluate was then subjected to cation-exchange chromatography using a MonoS 10/100 column (GE healthcare) and eluted with a linear gradient of NaCl. Fractions containing GRK5 were combined and run on a Superdex 200 10/300 GL column (GE healthcare). GRK5 was aliquoted, flash frozen, and stored at -80 °C until use.

### Arrestin expression and purification

The parent construct for β-arrestin 1 (βarr1) is the long splice variant of human, cysteine-free (C59V, C125S, C140L, C150V, C242V, C251V, C269S) β-arrestin 1. This construct is modified with an N-terminal 6x Histidine tag, followed by a 3C protease site, a GG linker, AviTag and GGSGGS linker. The sequence was codon-optimized for expression in *E. coli* and cloned into a pET-15b vector. Point mutations were prepared using site-directed mutagenesis. β-arrestin 1 (1-382) was prepared by truncating β-arrestin 1 at residue 382. All arrestin constructs used were prepared as follows: NiCo21(DE3) competent *E. coli* (NEB) were transformed, and large-scale cultures were grown in TB + ampicillin at 37°C until an OD_600_ of 1.0. Cells were then transferred to room temperature and induced with 25 μM IPTG when the OD_600_ reached 2.0. Cells were harvested 20 h post induction and resuspended in lysis buffer [50 mM Hepes pH 7.4, 500 mM NaCl, 15% glycerol, 7.13 mM β-mercaptoethanol (BME)] to a final volume of 40 mL/L of cells. Cells were lysed by sonication and the clarified lysate applied to nickel sepharose and batch incubated for 1.5h at 4 °C. The resin was washed with 10 column volumes of wash buffer (20 mM HEPES pH 7.4, 500 mM NaCl, 10% glycerol, 7.13 mM BME) + 20 mM imidazole, followed by 10 column volumes of wash buffer + 40 mM imidazole. The protein was then eluted with 5 column volumes of wash buffer + 200mM imidazole and dialyzed overnight in 100x volume of dialysis buffer (20 mM Hepes 7.4, 200 mM NaCl, 2 mM BME, 10% glycerol) in the presence of 1:10 (w:w) of 3C protease. The digested protein was then subjected to reverse-Nickel purification and diluted with dialysis buffer containing no NaCl to bring the NaCl concentration to 75mM. The protein was then purified by ion exchange chromatography (mono Q 10/100 GL, GE Healthcare), followed by SEC using a Superdex 200 increase 10/300 GL column (GE Healthcare) with SEC buffer (20 mM HEPES pH 7.4, 300 mM NaCl, 10% glycerol). Purified protein was concentrated to between 100- 300 μM using a 30 kDa spin concentrator and aliquots were flash-frozen in liquid nitrogen and stored at -80 °C until use.

### Arrestin labeling and biotinylation

Following SEC, elution peak fractions were pooled to a concentration of 10-20 μM and labeled with fluorophore(s): monobromobimane (mBBr), Thermo Fisher Scientific M1378; N,N’-Dimethyl- N-(Iodoacetyl)-N’-(7-Nitrobenz-2-Oxa-1,3-Diazol-4-yl)Ethylenediamine (IANBD amide), Thermo Fisher Scientific D2004; or a 1:3 mixture of Alexa Fluor 488 C5 Maleimide, Thermo Fisher Scientific A10254, and Atto647N Maleimide, ATTO TEC AD647N-41, respectively. Fluorophores were dissolved to in DMSO and added at 10x molar excess over protein, then allowed to react for 1 h at room temperature prior to quenching with cysteine (10x molar excess over fluorophore). The labeling reaction was further incubated for 10 minutes after cysteine addition, after which samples were spin filtered and subjected to a second round of size-exclusion chromatography, as detailed above, to remove free dye. The purified, was concentrated to between 100-300 μM using a 30 kDa spin concentrator and aliquots were flash-frozen in liquid nitrogen and stored at - 80 °C until use.

Arrestins (SEC-pure) were biotinylated using recombinant BirA enzyme, according to commercial protocols (Avidity), with exception that biotinylation was carried out for 12 h at 4 °C, rather than 30 °C. After biotinylation was complete, the reaction was flowed over 100 μL (packed) of nickel Sepharose, equilibrated in arrestin SEC buffer and supplemented with 10 mM imidazole, then washed with 200 μL of the equilibration buffer. The combined flow-through and wash fractions were then purified by size-exclusion as described above.

### NTSR1 phosphorylation

NTSR1 (2.5 μM) was equilibrated in phosphorylation buffer (20 mM bis-tris propane (BTP) pH 7.5, 35 mM NaCl, 5 mM MgCl2, 20 µM NTS_8-13_, 20 μM C8-PI(4,5)P2, 0.05 mM TCEP, 0.002% MNG, 0.0002% CHS) at 25 °C with gentle mixing for 1 h. GRK5 was added to the reaction to a final concentration of 200 nM, and briefly incubated while the reaction was warmed from 25 °C to 30 °C. ATP was added to a final concentration of 1 mM. Upon completion, the reaction was supplemented with CaCl_2_ to a final concentration of 2 mM and applied to an equilibrated M1 FLAG immunoaffinity resin and washed with buffer containing 0.004% LMNG, 0.004% CHS, 20 mM HEPES pH 7.4, 100 mM NaCl, 0.2 μM NTS_8-13_, 2 mM CaCl_2_. The receptor was eluted with buffer containing 100 mM NaCl, 20 mM HEPES pH 7.4, 0.004% LMNG, 0.004% CHS, 0.2 μM NTS_8-13_, 0.2 mg/mL 1x flag peptide (DYKDDDDK), 5 mM EDTA), followed by SEC using a Superdex 200 increase 10/300 GL column (GE Healthcare) with SEC buffer (20 mM HEPES pH 7.4, 100 mM NaCl, 0.004% LMNG, 0.0004% CHS).

### Analytical fluorescence-detection size-exclusion chromatography

In a final volume of 20 μL, NTSR1 (4.5 μM), the respective arrestin construct (9 μM), NTS_8-13_ peptide (50 μM) and diC8-PI(4,5)P2 (5 μM) were incubated in buffer containing 20 mM HEPES pH 7.4, 100 mM NaCl, 0.004% LMNG, 0.0004% CHS and 0.2 μM NTS_8-13_. Using a Prominence-i LC autosampler (Shimadzu), 10 μL was injected onto a ENrich size-exclusion chromatography 650 10 × 300 column (Bio-rad) pre-equilibrated in 20 mM HEPES pH 7.4 100 mM NaCl, 0.004 % LMNG, 0.004% CHS and 0.2 μM NTS_8-13_, and run at a flow rate of 0.8 ml/min. Tryptophan fluorescence was monitored at λ(EX) of 280 nm and λ(EM) of 340 nm. Peaks in the obtained size-exclusion chromatograms were modeled as gaussians, deconvolved and quantified (AUC) using Magic Plot 3 (Magic Plot).

### Surface plasmon resonance measurements

SPR experiments were performed using a GE Biacore T100 instrument. Approximately 300-400 resonance units (RU) of FPLC-purified biotinylated arrestin in HBS-P+ Buffer (GE Healthcare) were captured on an SA-chip (GE Healthcare), including a reference channel for online background subtraction of bulk solution refractive index and for evaluation of non-specific binding of analyte to the chip surface (Biacore T100 Control Software; GE Healthcare). All measurements were performed with 2-fold serial dilutions using 60 s association followed by a dissociation time of more than 240 s at 25 °C with a flow rate of at 30 μl min^−1^. Measurement of titrations at equilibrium were used to determine K_d_ values using Biacore Analysis Software (v.2.0.4, GE Healthcare) and fits to a total binding model were performed in GraphPad Prism 9. Regeneration was performed by 2 injections of 2 M MgCl_2_ for 10 s at 50 μl min^−1^ flow rate. In all cases regeneration resulted in a complete return to baseline. Single cycle measurements were performed as described above. All single cycle measurements were performed as triplicates and quantifications calculated to the RU_max_ of the individual immobilized ligands (arrestin proteins).

### Fluorescence anisotropy measurements

BODIPY-TMR phosphatidylinositol 4,5-bisphosphate (Echelon Biosciences) was dissolved to a stock concentration of 1 mM in 50 mM Hepes pH 7.4 and used at a final concentration of 4 nM in the assay. For the arrestin measurements, a two-fold dilution series of was made from a stock of βarr1 (1-382), yielding fourteen samples with final concentrations ranging from 150 µM to 0.02 µM. A control sample containing buffer only was included to measure the free anisotropy of BODIPY-PIP_2_. After mixing the BODIPY-PIP_2_ with arrestin or buffer, samples were incubated for 1h at room temperature prior to measurements. Samples were measured in five 20 µL replicates in a 384-well plate on a Tecan Infinite M1000 (Tecan Life Sciences), using an excitation wavelength of 530 nm, an emission wavelength of 573 nm and bandwidths of 5 nm. The obtained data was fit using to a one-site total binding model *Y* = *Bottom* + (*top* − *bottom*) /1 + 10*^HS^*^\**log*(*EC*50−*X*)^ where HS denotes the hill-slope.

### Bulk fluorescence measurements

Bulk fluorescence measurements were performed on either a Fluorolog instrument (Horiba) using FluorEssence v3.8 software and operating in photon-counting mode, or a Tecan Infinite M1000 PRO multimodal microplate reader (Tecan). Fluorolog measurements of bimane-labeled βarr1 constructs (NTSR1 experiments) were performed at final concentration of 0.4 μM [arrestin] in buffer containing 20 mM HEPES pH 7.4, 100 mM NaCl and 0.004% LMNG (w/v)/0.0004% CHS (w/v) supplemented with 4 μM NTS(8-13). For NTSR1 experiments the following concentrations were used: 4 μM NTSR1, 4.1 μM diC8-PI(4,5)P2, 50 μM V2Rpp (depending on condition). Samples were incubated for 1 h in the dark before measurement. Fluorescence data were collected in a quartz cuvette with 135 mL of sample. Bimane fluorescence was measured by excitation at 370 nm with excitation and emission bandwidth passes of 3 nm, and emission spectra were recorded from 400 to 550 nm in 2 nm increments with 0.1 s integration time. Care was taken to extensively rinse and argon-dry the cuvette between individual measurements. To remove background fluorescence, buffer spectra were collected using the same settings, and subtracted from each sample spectrum.

FRET measurements of AF488-AT647N-labeled βarr1 constructs were performed as described for bimane measurements, with the following differences: samples were excited at 476 nm with 3 nm excitation and 4 nm emission slit widths. Spectra were collected from 485 nm to 750 nm in 1 nm increments with 0.1 s integration time. FRET measurements in the absence of NTSR1 were performed in buffer containing 20 mM HEPES pH 7.4, 100 mM NaCl and 0.004% LMNG (w/v)/0.0004% CHS (no NTS). FRET measurements with NTSR1 were done with 0.5 μM NTSR1 and 0.5 μM diC8-PI(4,5)P2.

NBD spectra measured on the Tecan Infinite M1000 PRO were collected using 384- or 96-well (1/2 area) flat black Greiner plates with 50 or 100 μL of sample, respectively, at a final concentration of 0.5 μM βarr1 in buffer containing 20 mM HEPES pH 7.4, 100 mM NaCl and 0.004% LMNG (w/v)/0.0004% CHS. For NBD the following instrument settings were used: excitation: 490 nm, emission 510-580 nm (1 nm steps) with 20 s read time and 400 Hz flash mode.

Gain and z-position were optimized prior to reading. Bimane spectra were collected in white plates using the following instrument settings: excitation: 370 nm, emission 420-500 nm (1 nm steps) with 20 s read time and 400 Hz flash mode.

Efret values for FRET experiments were calculated as 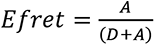 and normalized to donor intensity within a given experiment. Scaled FRET values (apo = 100, min(FRET) = 0) were fit to a single exponential decay function *Y* = (*Y*0 − *NS*) * *e*^−*K*\**x*^ + *NS* using the nls function in R for EC_50_ values (obtained as t_1/2_ for decay). NS denotes concentration-dependent non-specific signal. L167W-293NBD was fit using the same function. L68bim data was fit to a total binding model 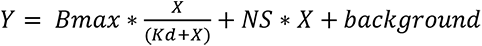, where background is a constant value. Fitting was independently performed both in R and with GraphPad Prism 9 for corroboration, values reported are from Prism 9.

---

**Figure S1.**
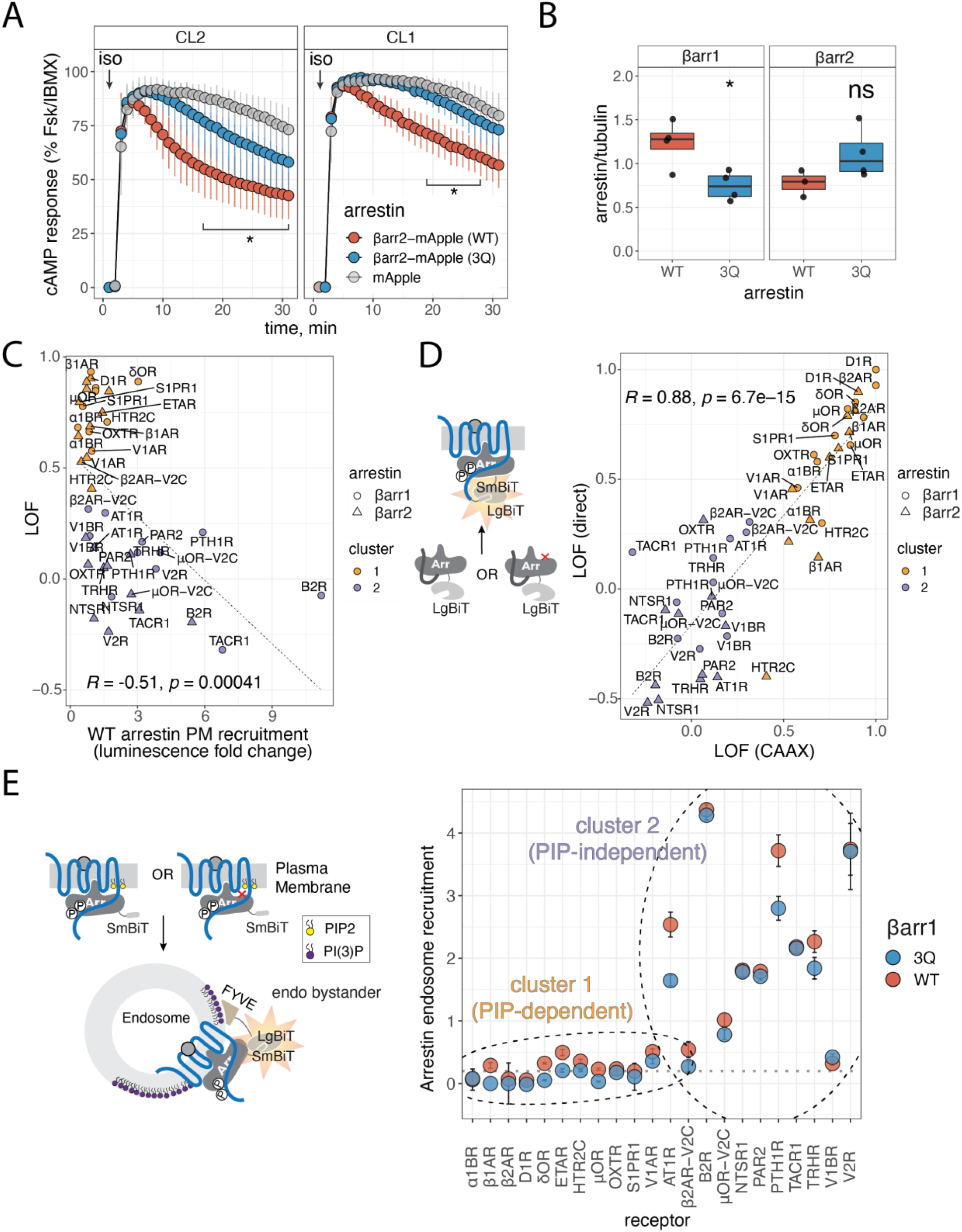
Arrestin phosphoinositide binding is required for plasma membrane recruitment to some GPCRs. A) cAMP response in HEK293 cells devoid of β-arrestins upon stimulation of endogenous β2AR with 100 nM isoproterenol (iso). Clone 1 (CL1) and Clone 2 (CL2) are independent βarr1/2 knock-out cell lines (O’Hare et al. 2017). Data are normalized to response with Forskolin (Fsk)/3-isobutyl-1-methylxanthine (IBMX) and show mean with 95% confidence intervals (*n*=3 independent experiments). Two-way analysis of variance (ANOVA), Tukey’s multiple comparison test. For CL2 * denotes p < 0.05 for WT vs. mApple over the interval of 17- 32 minutes, while 3Q vs. mApple was not significant. For CL1 * denotes p < 0.05 for WT vs. mApple over the interval of 19-29 minutes, while 3Q vs. mApple was not significant. B) Quantification of expression for βarr1 and βarr2 (both WT and 3Q) NanoBiT constructs, as determined by western blot (Supplementary Data Figure 2). Mean values of 3-4 independent experiments were compared by a two-tailed unpaired t-test, where ns denotes p > 0.05, * P ≤ 0.05. Boxplots: center line, median; box range, 25–75th percentiles; whiskers denote minimum– maximum values. Individual points are shown. C) LOF is only weakly correlated with recruitment of WT β-arrestins. Data are mean LOF and mean WT βarr1/2 recruitment. βarr1 recruitment is shown as circles and βarr2 recruitment is shown as triangles. Data are colored based on assigned cluster. Dashed line shows expected linear relationship and R is the Pearson coefficient, with - 0.51 reflecting a weak negative correlation. D) Plot of LOF data for plasma membrane bystander (CAAX) vs. LOF for direct recruitment. βarr1 recruitment is shown as circles and βarr2 recruitment is shown as triangles. Data are colored based on assigned cluster. Dashed line shows expected linear relationship and R is the Pearson coefficient, with 0.88 reflecting a very strong positive correlation. E) NanoBiT assay for measuring endosome translocation of βarr1. Cartoon of endosome bystander assay (left). βarr1 endosome recruitment data (right) with dashed ellipses to indicate clusters based on CAAX data. β-arrestin endosome recruitment determined by span of luminescence fold change. Data are mean ± SEM (*n*=3 independent experiments). Dashed line indicates three times the maximum signal measured in mock (receptor) transfected cells.

**Figure S2.**
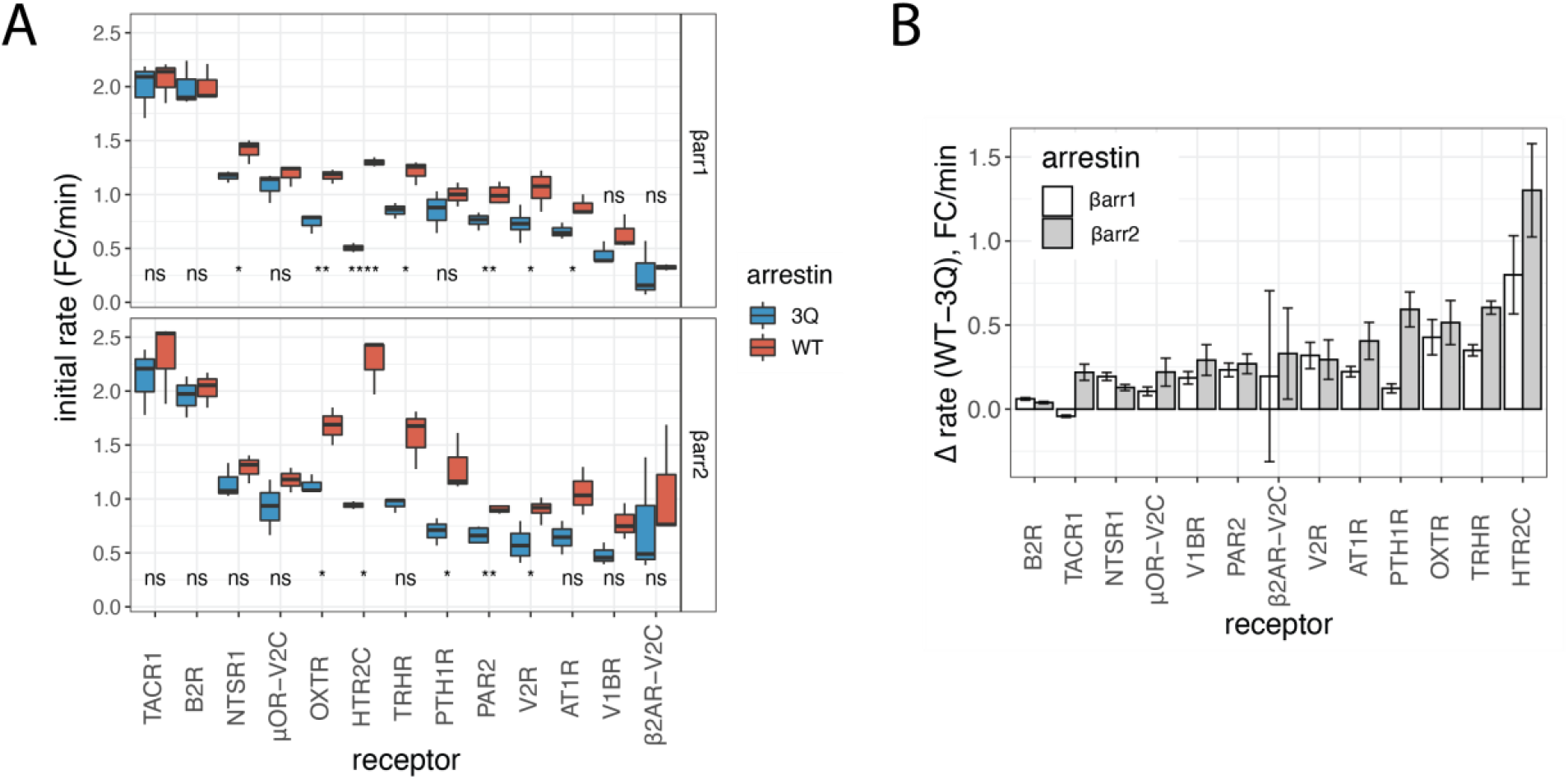
Loss of PIP binding slows β-arrestin recruitment to cluster 2 GPCRs. A) initial rate (0- 5 minutes post-agonist stimulation) expressed as luminescence fold-change (FC)/min. Data from *n*=3 independent experiments fit independently (see methods). Boxplots: center line, median; box range, 25–75th percentiles; whiskers denote minimum–maximum values. For each receptor, and for each βarr1 and βarr2 WT and 3Q were compared by a two-tailed unpaired t-test, where ns denotes p > 0.05, * P ≤ 0.05, ** P ≤ 0.01, **** P ≤ 0.0001. B) Data from A) expressed as a difference in rate shows that with the exception of βarr1-TACR1 all cluster 2 receptors show faster recruitment of WT β-arrestin1/2 than corresponding 3Q mutant. Data are mean ± SEM (*n*=3 independent experiments).

**Figure S3.**
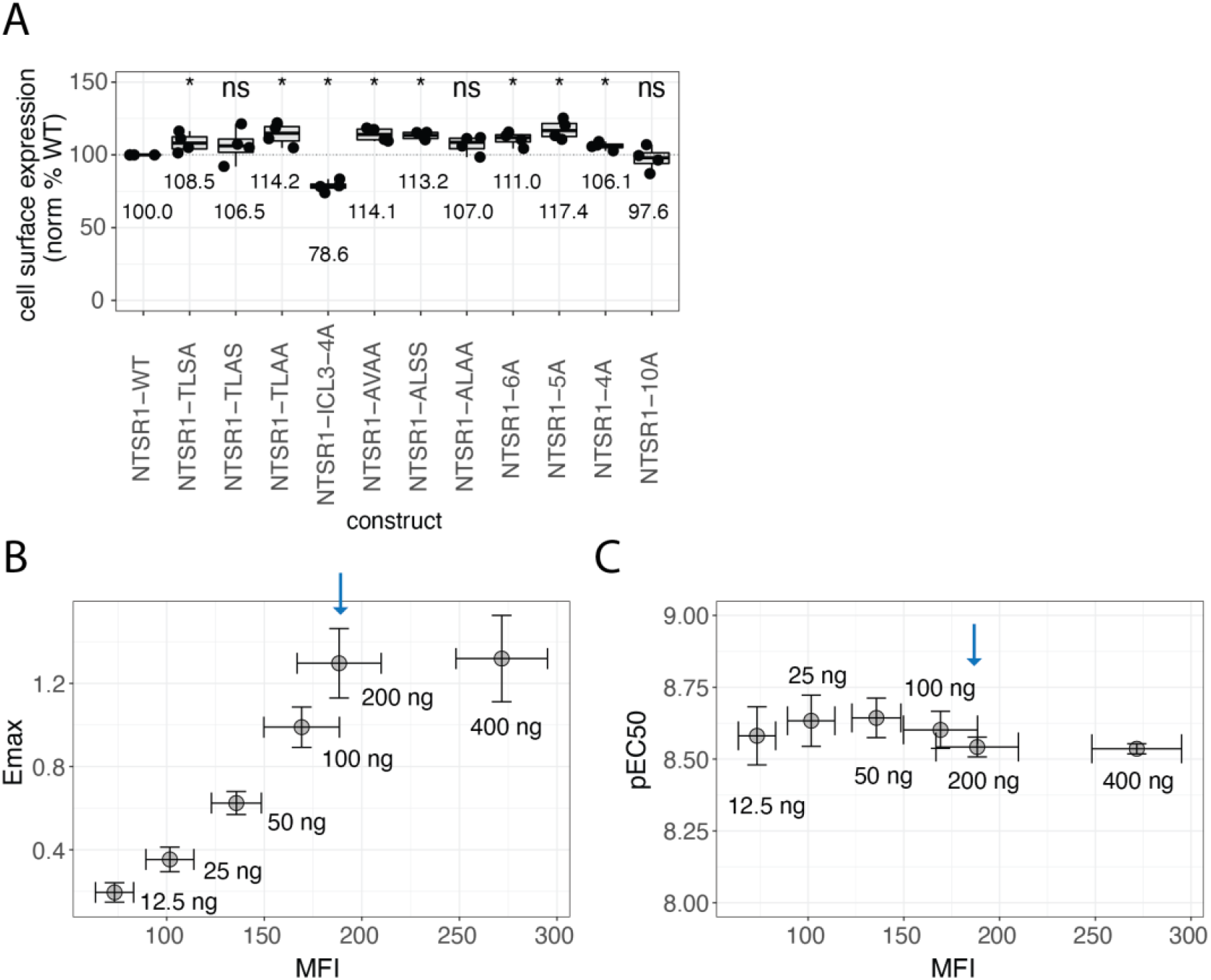
Arrestin recruitment to NTSR1 mutants can be measured by NanoBiT recruitment assay. A) Expression of NTSR1 constructs in HEK293A cells used for NanoBiT assays. Boxplots: center line, median; box range, 25–75th percentiles; whiskers denote minimum–maximum values. Individual points are shown. Values are mean, relative to NTSR1-WT (*n*=4 independent experiments). For each construct, a comparison to NTSR1-WT by a two-tailed unpaired Wilcoxon test was performed, where ns denotes p > 0.05, * P ≤ 0.05. B) Direct complementation NanoBiT assay Emax for Sm-βarr1 interaction with Lg-CAAX for cells expressing NTSR1-WT as a function of mean fluorescence intensity (MFI), as determined by cell-surface staining. Amount of NTSR1- WT DNA transfected is written; blue arrow denotes 200 ng, the amount used in recruitment assays in Figure 2. C) As B, except the pEC_50_ of recruitment response upon NTS stimulation is plotted vs. MFI, instead of Emax. In both B and C, points represent mean values and error bars indicate 95% CI (*n*=3 independent experiments).

**Figure S4.**
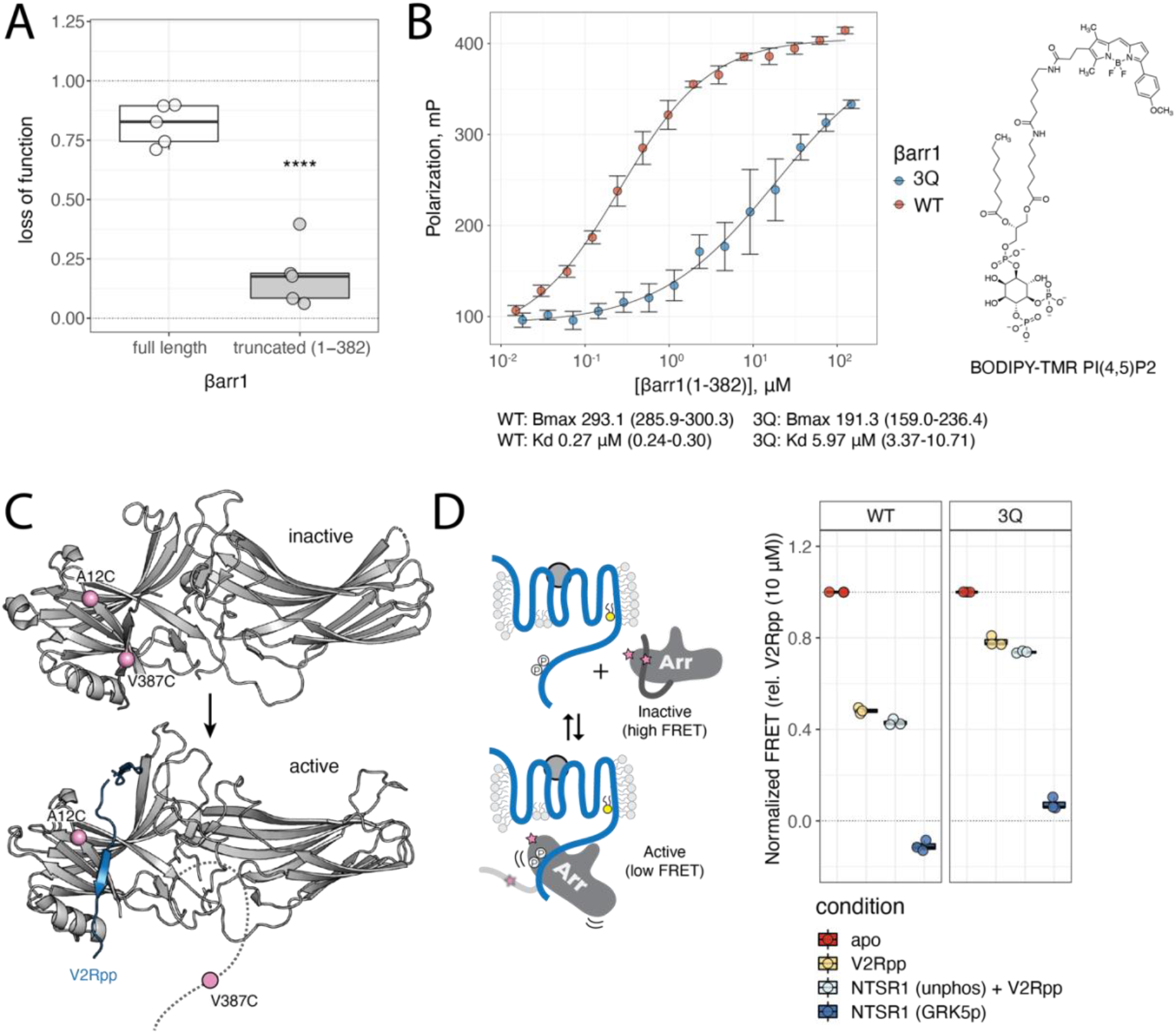
PIP binding stabilizes core-engaged arrestin complexes. A) LOF in complexing efficiency as determined by SEC. LOF = 1 corresponds to complete loss of complex formation, while LOF = 0 corresponds to no difference in complexing efficiency between WT and 3Q βarr1 (*n*=5 independent experiments). Boxplots: center line, median; box range, 25–75th percentiles; whiskers denote minimum–maximum values. Individual points are shown. compared by a two-tailed unpaired t-test, where **** P ≤ 0.0001. B) Binding of BODIPY-TMR PI(4,5)P2 to βarr1 (1-382) protein (WT or 3Q). Points are mean and error bars reflect 95% CI (*n*=5 independent experiments). Data were fit to a logistical function as described in methods and best fit values for B_max_ and K_d_ are provided with 95% CI in parentheses. C) Structure of transition from inactive (PDB: 1G4M) to active (PDB: 4JQI) βarr1 involves displacement of the βarr1 C-terminus (dark grey) by V2Rpp (blue). Two cysteine residues were added to a cys-less βarr1 background at positions A12 and V387 (pink spheres). These positions were labeled with fluorophores that, through FRET, allow for monitoring the position of the C-terminus. D) When labeled with a FRET pair, βarr1-12C/387C shows a high-FRET state in the absence of V2Rpp, and a low-FRET state when the βarr1 C-terminus is displaced by V2Rpp. FRET measured when βarr1 (WT or 3Q)- 12C/387C-AF488-AT647N is bound to V2Rpp (0.5 μM), NTSR1 (GRK5p, 0.5 μM), or NTSR1 (unphosphorylated, 0.5 μM)+V2Rpp (0.5 μM). All samples containing NTSR1 were supplemented with diC8-PI(4,5)P2 (0.5 μM). Apo βarr1 (WT or 3Q)-12C/387C-AF488-AT647N was normalized to 1.0 and βarr1 (WT or 3Q)-12C/387C-AF488-AT647N + V2Rpp (10 μM) was normalized 0.0 for each experiment (*n*=3 independent measurements) (right). Boxplots: center line, median; box range, 25–75th percentiles; whiskers denote minimum–maximum values. Individual points are shown.

**Figure S5.**
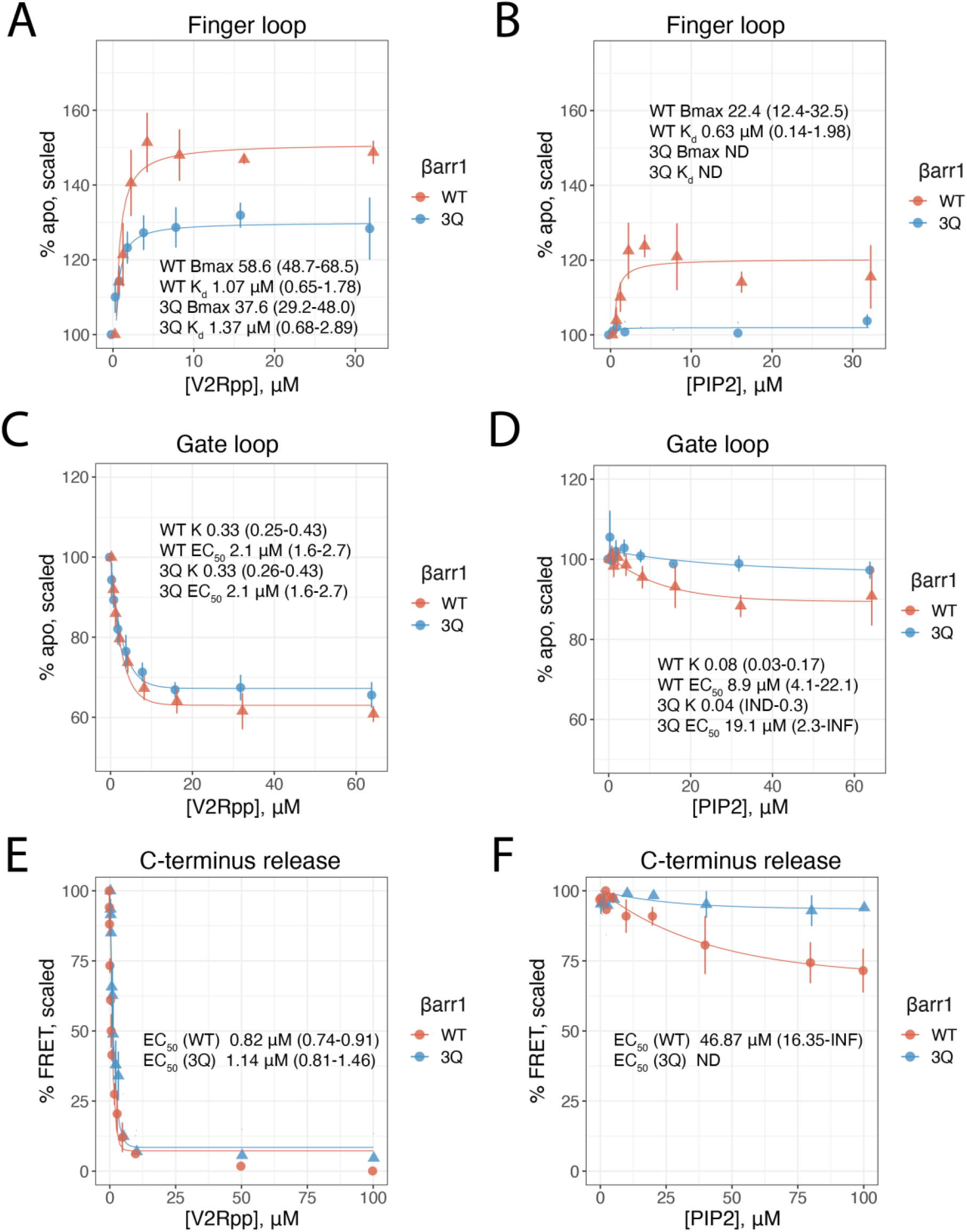
PIP_2_ allosterically triggers movement of the arrestin C-tail, but not release. A-B) Finger loop (L68C-bim) responses. %apo is scaled such that the fluorescence intensity (at λmax) for apo arrestin is 100% and each condition is scaled as a factor of apo. ND denotes not determined values. Values for B_max_ (max response) and K_d_ (based on single-site binding fitting) are provided and ranges in parentheses correspond to 95% CI. Points are mean and error bars reflect 95% CI (*n*=3 independent experiments). C-D) Gate loop (L167W-293C-NBD) responses. %apo is scaled such that the fluorescence intensity (at λmax) for apo arrestin is 100% and each condition is scaled as a factor of apo. Values for EC_50_ (half maximal response) and k (rate constant based on single exponential decay) are provided and ranges in parentheses correspond to 95% CI. Points are mean and error bars reflect 95% CI (*n*=3 independent experiments). E-F) C-terminus release (A12C-V387C-AF488-AT647N) responses. %FRET is scaled such that apo arrestin is 100% and the highest concentration of V2Rpp (100 uM) is 0%. INF denotes infinite upper bound. ND denotes not determined values. Range of EC_50_ values is indicated in parentheses and represents 95% CI. Points represent mean and error bars reflect 95% CI (*n*=3 independent experiments).

**Figure S6.**
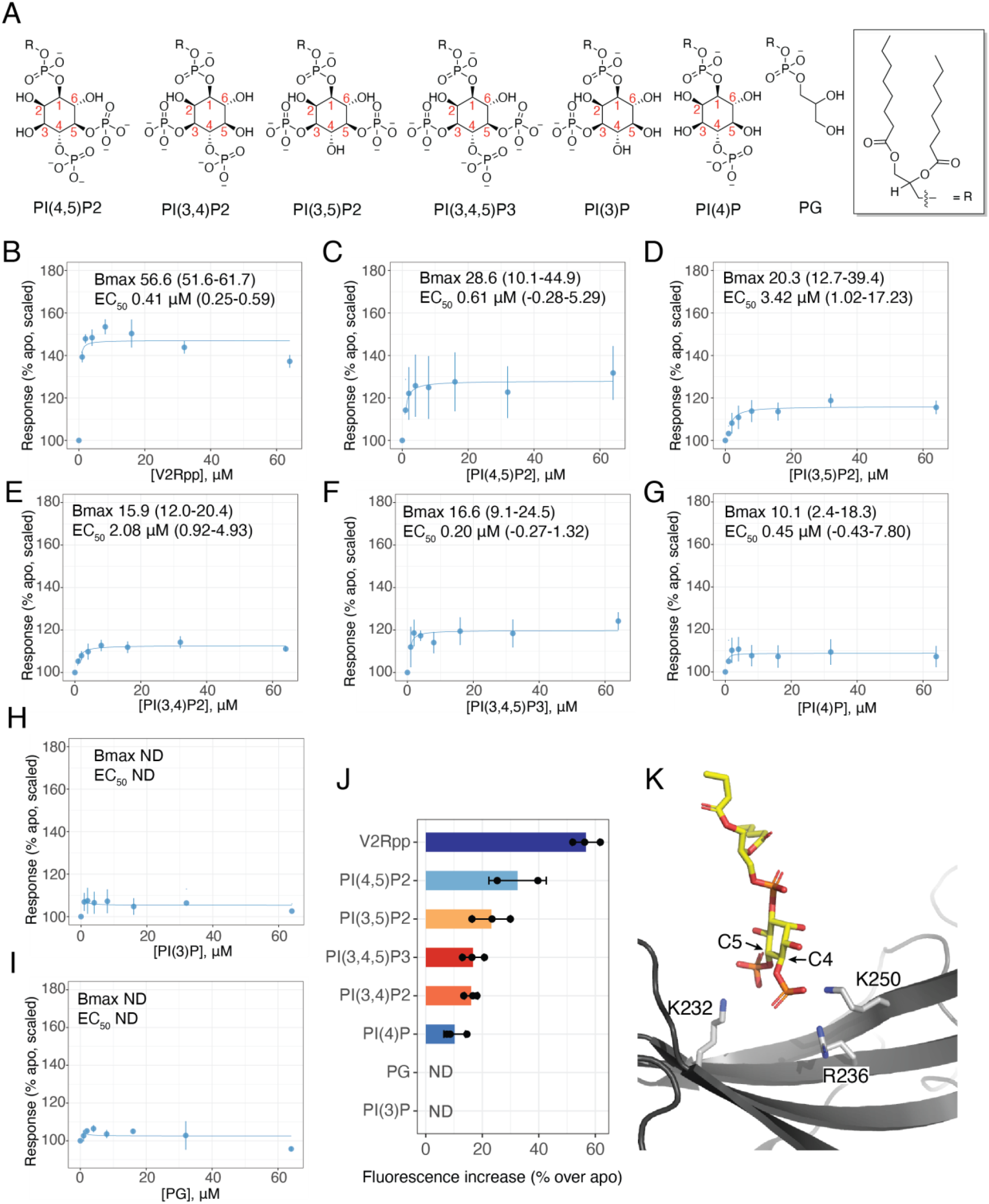
Plasma membrane PIPs promote conformational changes in arrestin. A) Structure of soluble lipid derivatives examined in this work. B-I) Concentration response curves for L68bim βarr1. Values for B_max_ (max response) and EC_50_ (based on single-site binding fitting) are provided and ranges in parentheses correspond to 95% CI. Points are mean and error bars reflect 95% CI (*n*=3 independent experiments). ND denotes not determined values. J) Summary of effect size (B_max_) for different lipids with L68bim βarr1. Values represent B_max_ obtained from fitting independent experiments. ND is used for PG and PI(3)P as data could not be fit. Bars represent mean B_max_ and error bars denote standard deviation across the fits. K) A PI(4,5)P2 derivative bound in the C-terminal domain of βarr1 (PDB: 6UP7). The side chain of K232 was modeled based on PDB: 4JQI as it was not ordered in PDB: 6UP7.

**Figure S7.**
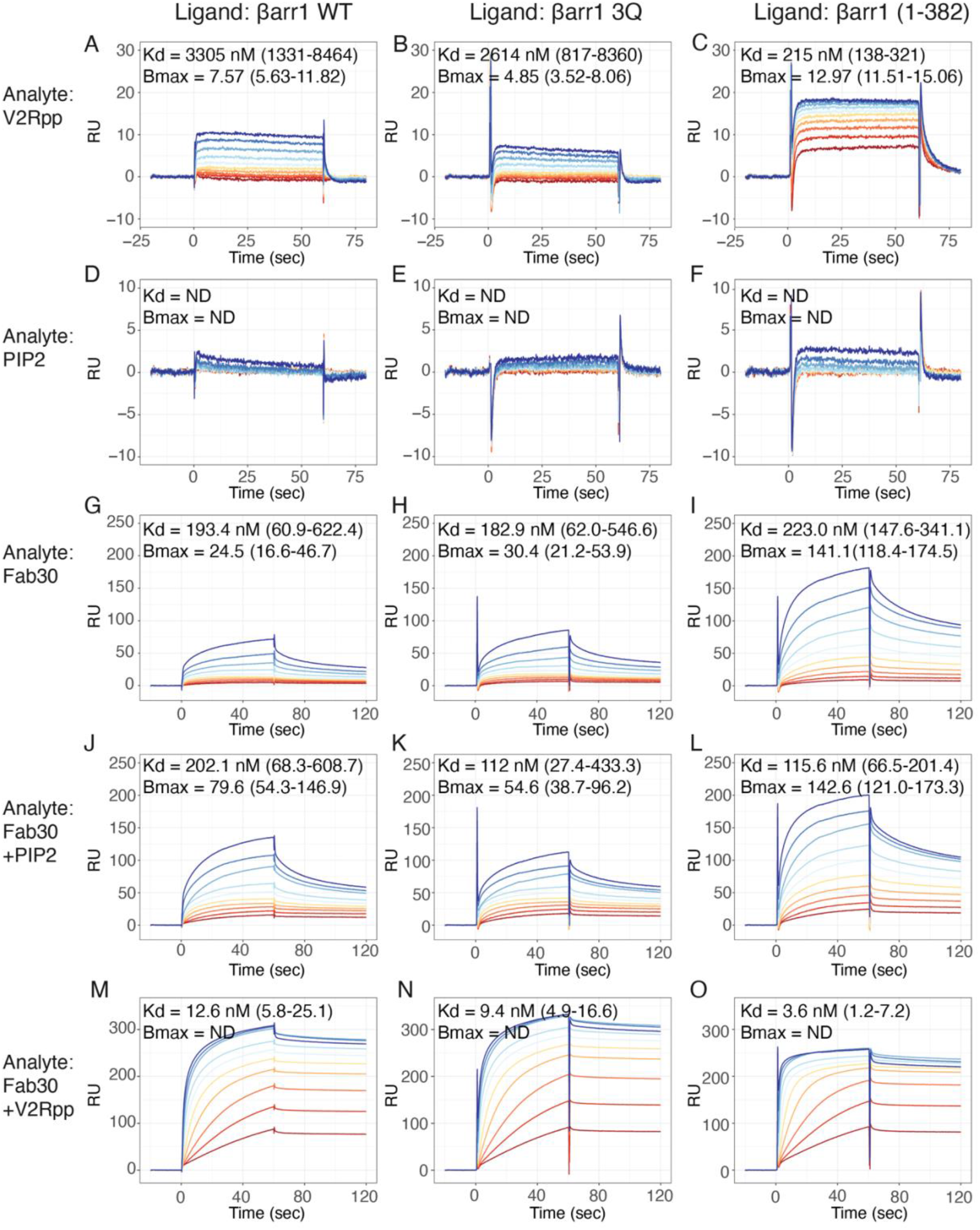
Titration of interactions with immobilized βarr1 by SPR. Immobilized βarr1 construct (ligand) is listed as column heading. Rows represent analyte flowed during titration. K_d_ and B_max_ values obtained as described in methods are listed with 95% CI range in parentheses. ND denotes value not determined. PIP_2_ binding was not fit (D-F), whereas B_max_ for Fab30+V2Rpp (M-O) failed to converge. In all cases the darkest blue curve corresponds to the highest concentration and the darkest red curve corresponds to the lowest concentration. Titrations, as described in methods, for V2Rpp and PIP_2_ (A-F) ranged from 40 μM to 78.1 nM. Titrations of Fab30 ranged from 2000 nM to 3.9 nM with fixed concentration of V2Rpp (40 μM) or PIP_2_ (40 μM), as indicated. Vertical axes are raw RU, and not corrected for channel loading.

