## Supplementary Data 1-3 for "Membrane phosphoinositides stabilize GPCR-arrestin complexes and provide temporal control of complex assembly and dynamics"

| GPCR | Abbreviation | Ligand | Log concentrations tested |
|---|---|---|---|
| Tachykinin Receptor 1 | TACR1 | Substance P | -15, -11, -10, -9, -8, -7, -6, -5 |
| B2 bradykinin receptor | B2R | Bradykinin | -15, -11, -10, -9, -8, -7, -6, -5 |
| Neurotensin receptor type 1 | NTSR1 | Neurotensin | -15, -10, -9, -8, -7, -6, -5.5, -5 |
| Vasopressin V2 receptor | V2R | AVP | -15, -9, -8.5, -8, -7.5, -7, -6.5, -6 |
| Proteinase-activated receptor 2 | PAR2 | PAR2 peptide | -15, -7.5, -7, -6.5, -6, -5.5, -5, -4.5 |
| Mu-type opioid receptor V2R CT chimera | μOR-V2C | DAMGO | -15, -9, -8, -7, -6, -5, -4.5, -4 |
| Thyrotropin-releasing hormone receptor | TRHR | TRH | -15, -10, -9, -8, -7, -6, -5, -4 |
| Parathyroid hormone/parathyroid hormone-related peptide receptor | PTH1R | PTH | -15, -10, -9, -8, -7, -6.5, -6, -5.5 |
| Vasopressin V1b receptor | V1BR | AVP | -15, -9, -8.5, -8, -7.5, -7, -6.5, -6 |
| Type-1 angiotensin II receptor | AT1R | AngII | -15, -11, -10, -9, -8, -7, -6, -5 |
| Oxytocin receptor | OXTR | Oxytocin | -15, -10, -9, -8, -7, -6, -5.5, -5 |
| Beta-2 adrenergic receptor V2R CT chimera | β2AR-V2C | Isoproterenol | -15, -9, -8, -7, -6, -5, -4, -3.5 |
| 5-hydroxytryptamine receptor 2C | HTR2C | Serotonin | -15, -9, -8, -7, -6, -5, -4, -3.5 |
| Vasopressin V1a receptor | V1AR | AVP | -15, -9, -8.5, -8, -7.5, -7, -6.5, -6 |
| Alpha-1B adrenergic receptor | α1BR | Norepinephrine | -15, -9, -8, -7, -6, -5, -4.5, -4 |
| Beta-1 adrenergic receptor | β1AR | Isoproterenol | -15, -9, -8, -7, -6, -5, -4, -3.5 |
| Endothelin receptor type A | ETAR | Endothelin | -15, -11, -10, -9, -8, -7, -6.5, -6 |
| Sphingosine 1-phosphate receptor 1 | S1PR1 | S1P | -15, -10, -9, -8, -7, -6.5, -6, -5.5 |
| Mu-type opioid receptor | μOR | DAMGO | -15, -9, -8, -7, -6, -5, -4.5, -4 |
| Delta-type opioid receptor | δOR | Met-Enk | -15, -10, -9, -8, -7, -6, -5.5, -5 |
| Beta-2 adrenergic receptor | β2AR | Isoproterenol | -15, -9, -8, -7, -6, -5, -4, -3.5 |
| D(1A) dopamine receptor | D1R | Dopamine | -15, -7, -6, -5.5, -5, -4.5, -4, -3.5 |

**Supplementary Table 1.** GPCR ligands and concentrations used in cell experiments. Vehicle treatment is denoted by a log(conc) of -15 for purposes of fitting.

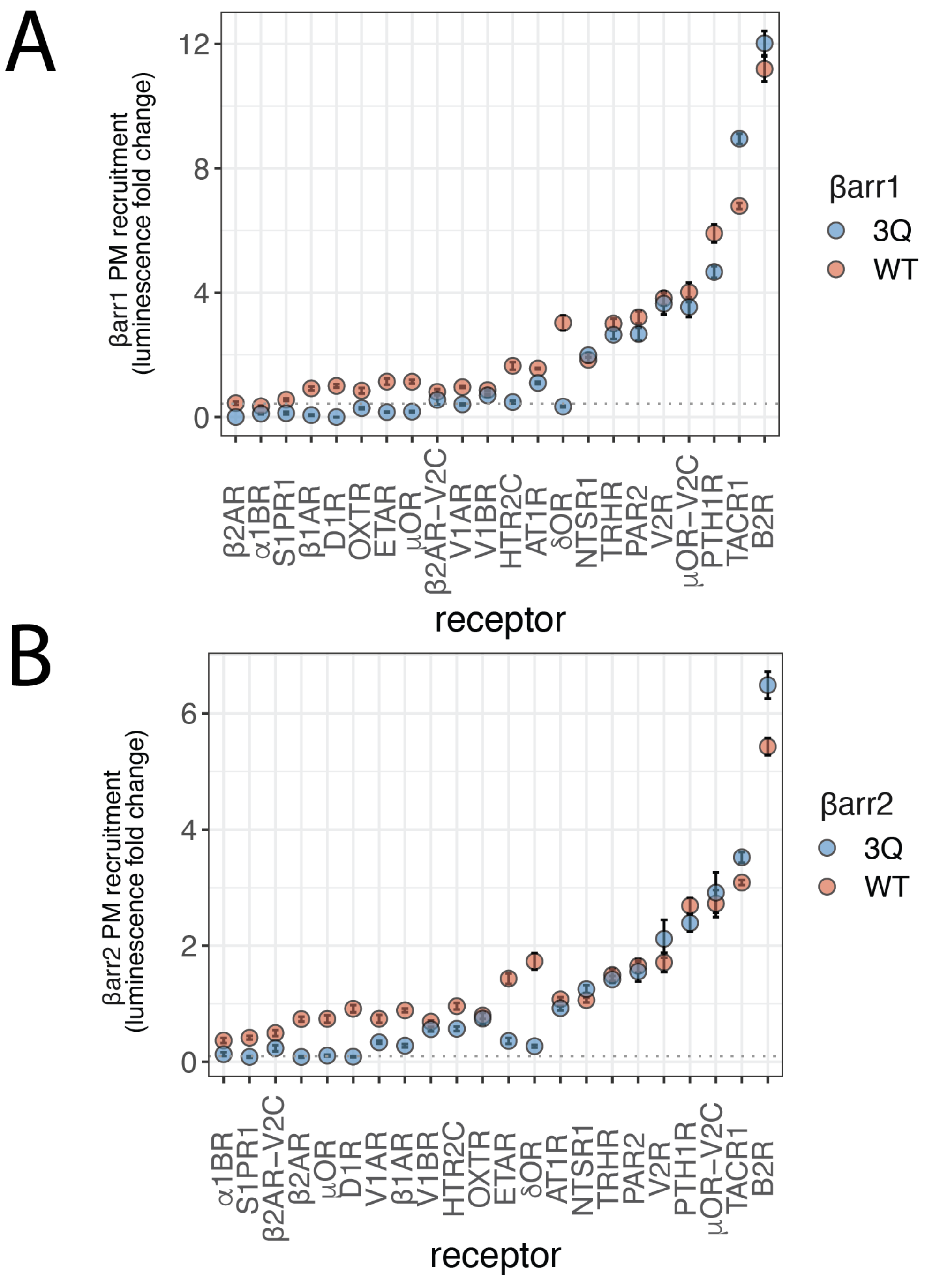


**Supplementary Data Figure 1.** Recruitment amplitude for arrestins to a CAAX plasma membrane bystander using NanoBiT assay. Points reflect best fit values from three independent experiments and error bars show standard error of fits. Vertical axis is scaled such that Luminescence fold-change refers to the span of the fit (top – bottom) A) βarr1 data. Dashed horizontal line corresponds to three times the max signal measured with receptor mock-transfected cells y = 0.43; B) βarr2 data. Dashed horizontal line corresponds to three times the max signal measured with receptor mock-transfected cells y = 0.096.

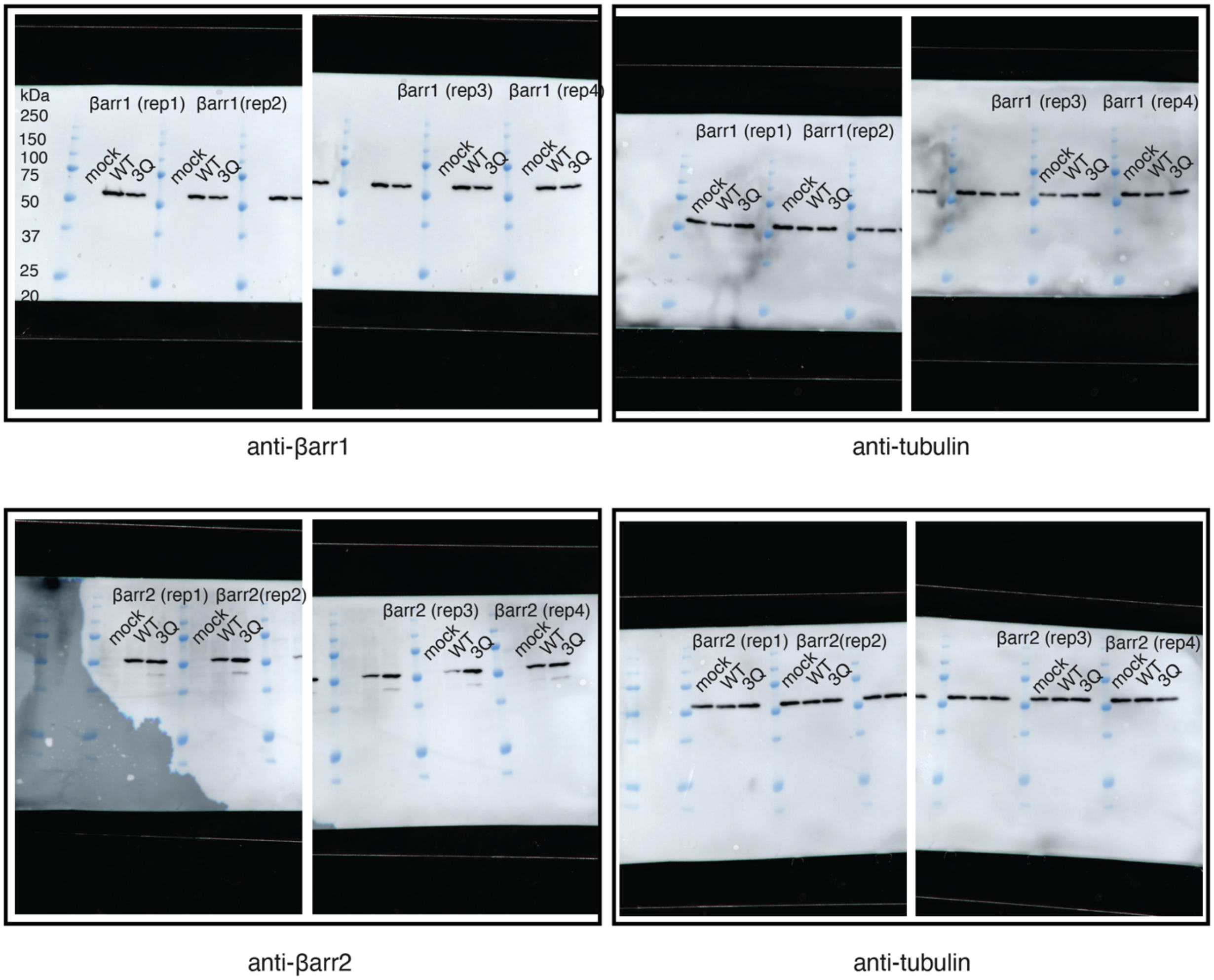


**Supplementary Data Figure 2.** Full western blots of βarr1/2-SmBiT both WT and 3Q for measuring expression levels. Representative protein marker labeled top left.

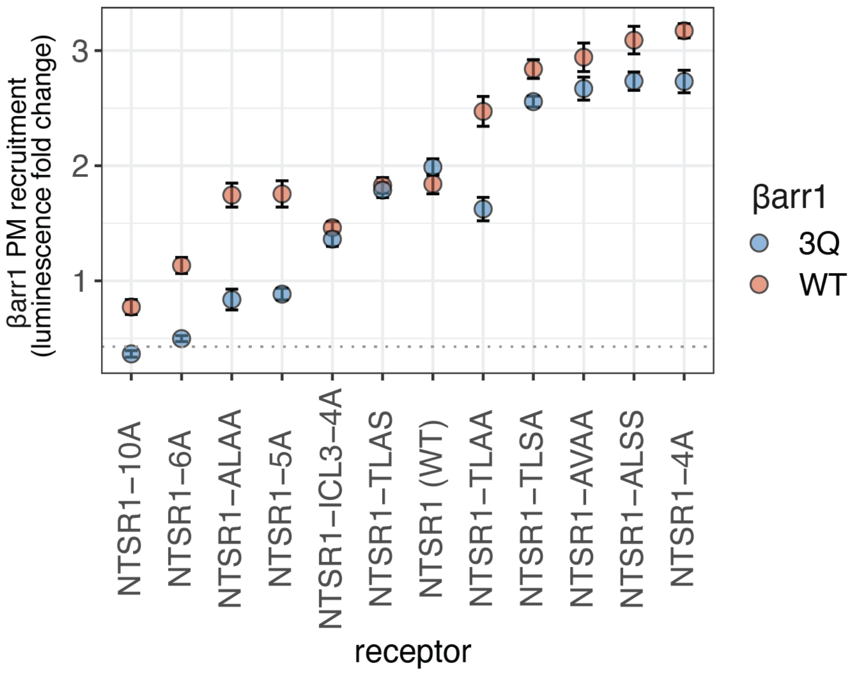


**Supplementary Data Figure 3.** NanoBiT recruitment amplitudes for βarr1 recruitment NTSR1 constructs using a CAAX plasma membrane bystander. Points reflect best fit values from three independent experiments and error bars show standard error of fits. Vertical axis is scaled such that luminescence fold-change refers to the span of the fit (top – bottom). Dashed horizontal line corresponds to three times the max signal measured with receptor mock-transfected cells y = 0.43.

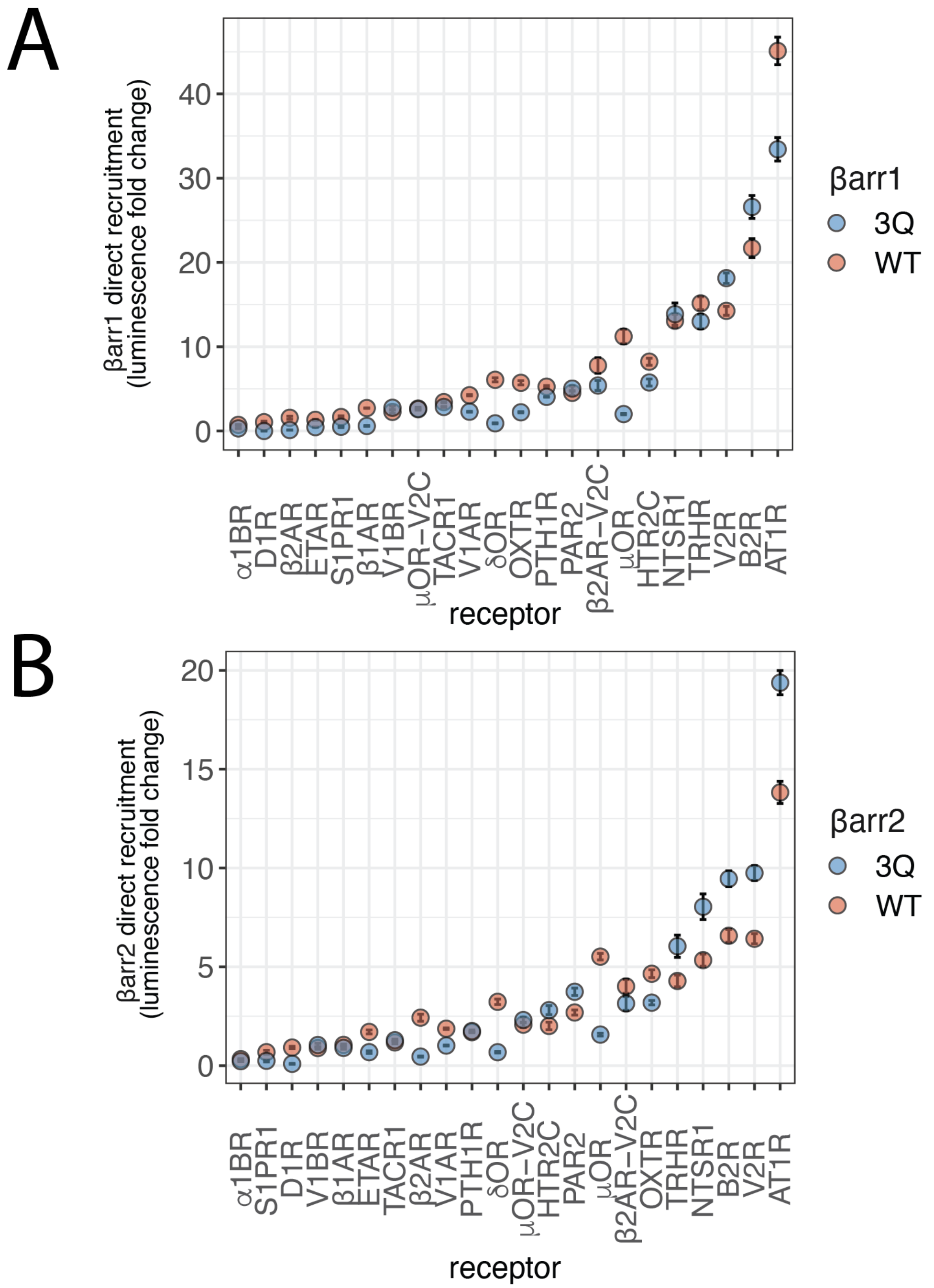


**Supplementary Data Figure 4.** Direct recruitment assay for LgBiT labeled arrestins to SmBiT tagged GPCRs. Points reflect best fit values from three independent experiments and error bars show standard error of fits. Vertical axis is scaled such that luminescence fold-change refers to the span of the fit (top – bottom) A) βarr1 data; B) βarr2 data.

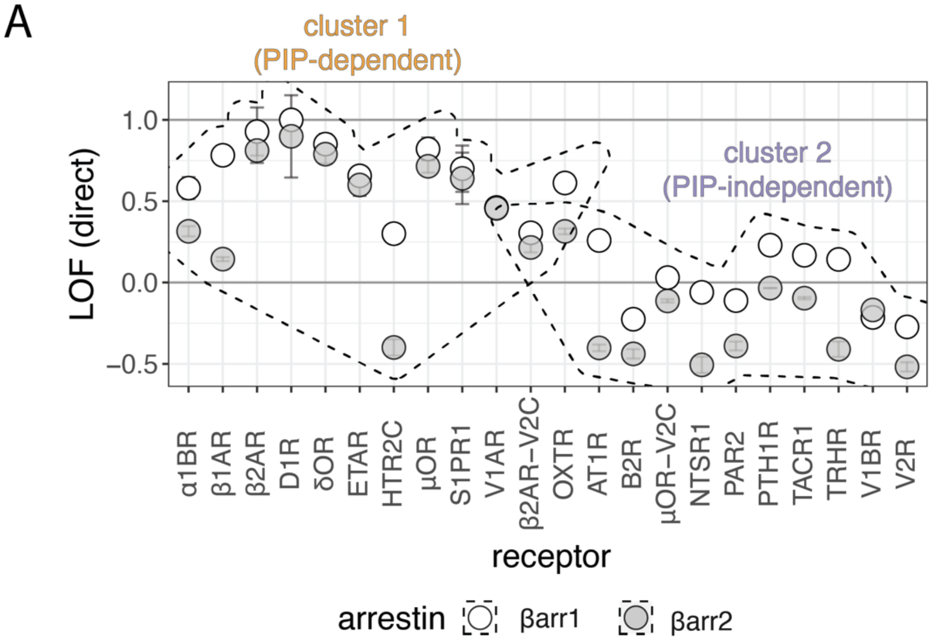


**Supplementary Data Figure 5.** A) LOF index measured by direct recruitment NanoBiT assay shows 3Q-dependence on arrestin recruitment. Data are mean ± SEM (n=3 independent experiments). Dashed shapes group cluster 1 and cluster 2 receptors. Both V1AR (βarr1 and βarr2 points overlap) pairs belong to cluster 1. β2AR-V2C βarr1 and OXTR βarr2 recruitment belong to cluster 2.

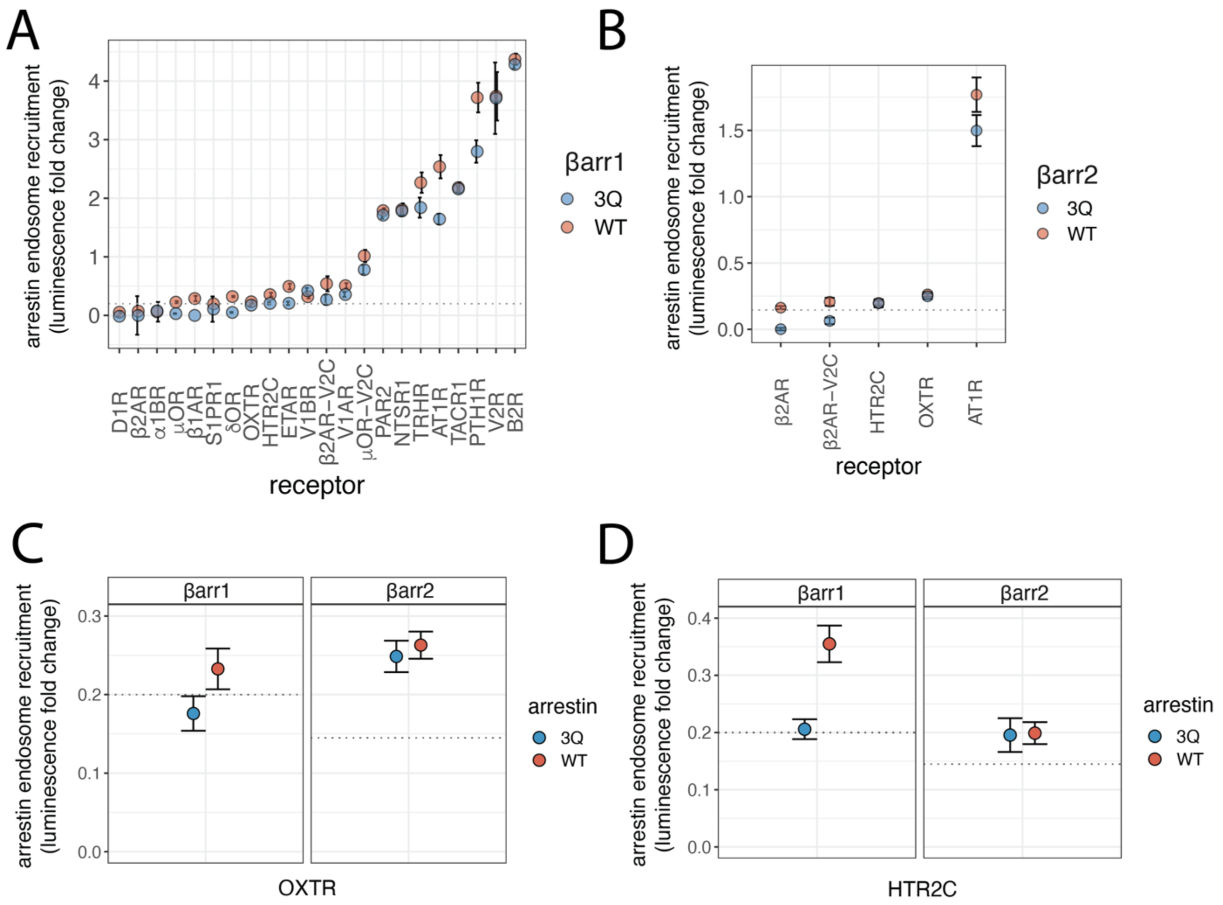


**Supplementary Data Figure 6.** Recruitment amplitude for arrestins to endosomes as measured by an endosome bystander NanoBiT assay. Points reflect best fit values from three independent experiments and error bars show standard error of fits. Vertical axis is scaled such that luminescence fold-change refers to the span of the fit (top – bottom) A) βarr1 data: dashed horizontal line corresponds to three times the max signal measured with receptor mock-transfected cells y = 0.20; B) βarr2 data, for selected receptors: dashed horizontal line corresponds to three times the max signal measured with receptor mock-transfected cells y = 0.14. C-D) Zooms of arrestin translocation to endosomes measured by endosome bystander NanoBiT assay for cells expressing OXTR and HTR2C, respectively. Dashed horizontal lines corresponds to three times the max signal measured with receptor mock-transfected cells, as stated in B) and C).

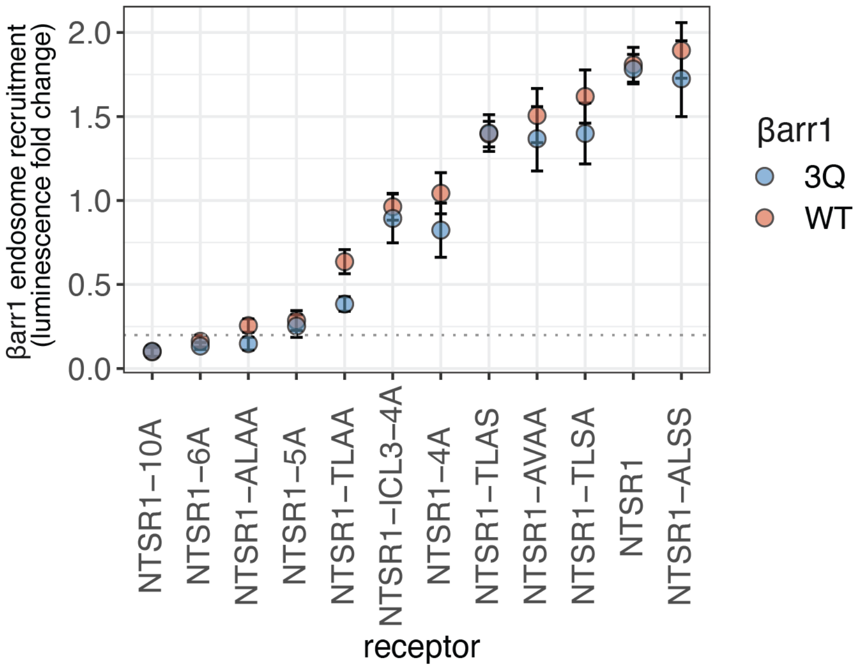


**Supplementary Data Figure 7.** NanoBiT recruitment amplitudes for βarr1 recruitment to endosomes for cells expressing NTSR1 constructs. Points reflect best fit values from three biological replicates and error bars show standard error of fits. Vertical axis is scaled such that luminescence fold-change refers to the span of the fit (top – bottom). Dashed horizontal line corresponds to three times the max signal measured with receptor mock-transfected cells y = 0.20.

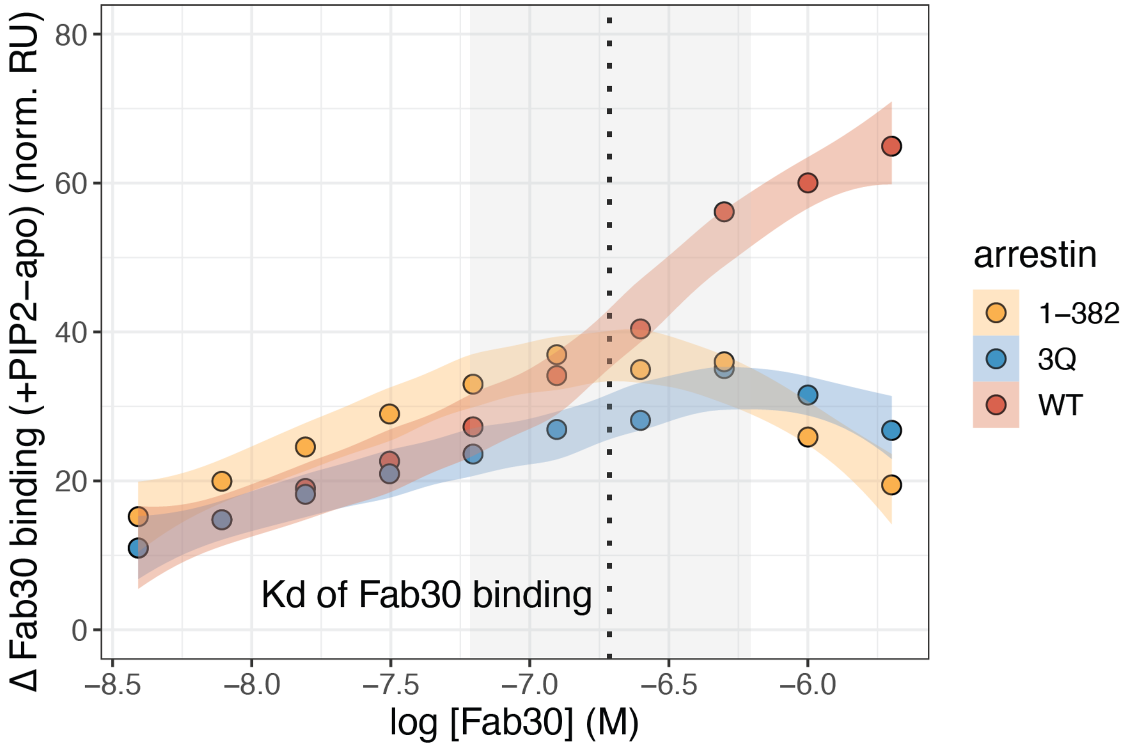


**Supplementary Data Figure 8.** Effect of $PIP_2$ on Fab30 binding by SPR. Points indicate steady-state binding of Fab30 to βarr1, at varying concentrations of Fab30. Titrations were performed in the presence (+$PIP_2$) or absence (apo) of $PIP_2$, as shown in Fig S7 panels G-L. Δ Fab30 binding was determined by subtracting the response for apo from that with fixed saturating PI(4,5)P2 (40 μM). RU values are normalized based on the immobilized RU of each arrestin. Vertical dashed line denotes best fit Kd value for Fab30 binding WT βarr1, and the grey box indicates the 95% CI (from Fig S7G). For each WT, 3Q and 1-382 βarr1 a LOWESS smoothing is used to join the points and standard error is indicated by the colored range. Data are from a single independent experiment.

| Figure | Type | Ligand (bio+) | Ligand RU | Analyte 1 | Analyte 1 [nM] | Analyte 2 | Analyte 2 [nM] | Dilution Series (anal | # Dilutions | Injection Time (sec) | Injection Rate (ul/min) | Time Dissociation (sec) | Regeneration | Regen Rate (ul/min) |
|---|---|---|---|---|---|---|---|---|---|---|---|---|---|---|
| Fig Suppl 7 | Titration | barr1 WT | 327 | Fab30 | 2000 | | | 2 | 10 | 60 | 30 | 240 | 3 x 10 sec; 2M MgCl2 | 50 |
| Fig Suppl 7 | Titration | barr1 WT | 327 | PIP2 | 40000 | | | 2 | 10 | 60 | 30 | 240 | 3 x 10 sec; 2M MgCl2 | 50 |
| Fig Suppl 7 | Titration | barr1 WT | 327 | V2Rpp | 40000 | | | 2 | 10 | 60 | 30 | 240 | 3 x 10 sec; 2M MgCl2 | 50 |
| Fig Suppl 7 | Titration | barr1 WT | 327 | Fab30 | 2000 | PIP2 | 40000 | 2 | 10 | 60 | 30 | 240 | 3 x 10 sec; 2M MgCl2 | 50 |
| Fig Suppl 7 | Titration | barr1 WT | 327 | Fab30 | 2000 | V2Rpp | 40000 | 2 | 10 | 60 | 30 | 240 | 3 x 10 sec; 2M MgCl2 | 50 |
| Fig Suppl 7 | Titration | barr1 3Q | 336 | Fab30 | 2000 | | | 2 | 10 | 60 | 30 | 240 | 3 x 10 sec; 2M MgCl2 | 50 |
| Fig Suppl 7 | Titration | barr1 3Q | 336 | PIP2 | 40000 | | | 2 | 10 | 60 | 30 | 240 | 3 x 10 sec; 2M MgCl2 | 50 |
| Fig Suppl 7 | Titration | barr1 3Q | 336 | V2Rpp | 40000 | | | 2 | 10 | 60 | 30 | 240 | 3 x 10 sec; 2M MgCl2 | 50 |
| Fig Suppl 7 | Titration | barr1 3Q | 336 | Fab30 | 2000 | PIP2 | 40000 | 2 | 10 | 60 | 30 | 240 | 3 x 10 sec; 2M MgCl2 | 50 |
| Fig Suppl 7 | Titration | barr1 3Q | 336 | Fab30 | 2000 | V2Rpp | 40000 | 2 | 10 | 60 | 30 | 240 | 3 x 10 sec; 2M MgCl2 | 50 |
| Fig Suppl 7 | Titration | barr1 1-382 | 332 | Fab30 | 2000 | | | 2 | 10 | 60 | 30 | 240 | 3 x 10 sec; 2M MgCl2 | 50 |
| Fig Suppl 7 | Titration | barr1 1-382 | 332 | PIP2 | 40000 | | | 2 | 10 | 60 | 30 | 240 | 3 x 10 sec; 2M MgCl2 | 50 |
| Fig Suppl 7 | Titration | barr1 1-382 | 332 | V2Rpp | 40000 | | | 2 | 10 | 60 | 30 | 240 | 3 x 10 sec; 2M MgCl2 | 50 |
| Fig Suppl 7 | Titration | barr1 1-382 | 332 | Fab30 | 2000 | PIP2 | 40000 | 2 | 10 | 60 | 30 | 240 | 3 x 10 sec; 2M MgCl2 | 50 |
| Fig Suppl 7 | Titration | barr1 1-382 | 332 | Fab30 | 2000 | V2Rpp | 40000 | 2 | 10 | 60 | 30 | 240 | 3 x 10 sec; 2M MgCl2 | 50 |

**Supplemantary Data Table 2**

| Figure | Type | Ligand (bio+) | Ligand RU | Analyte 1 | Analyte 1 [nM] | Analyte 2 | Analyte 2 [nM] | # Replicates | Injection Time (sec) | Injection Rate (ul/min) | Time Dissociation (sec) | Regeneration | Regen Rate (ul/min) |
|---|---|---|---|---|---|---|---|---|---|---|---|---|---|
| Fig 5C | Single Cycle | barr1 WT | 411 | Fab30 | 1000 | | | 3 | 60 | 30 | 240 | 3 x 10 sec; 2M MgCl2 | 50 |
| Fig 5C | Single Cycle | barr1 WT | 411 | PIP2 | 40000 | | | 3 | 60 | 30 | 240 | 3 x 10 sec; 2M MgCl2 | 50 |
| Fig 5C | Single Cycle | barr1 WT | 411 | PI(3)P | 40000 | | | 3 | 60 | 30 | 240 | 3 x 10 sec; 2M MgCl2 | 50 |
| Fig 5C | Single Cycle | barr1 WT | 411 | PG | 40000 | | | 3 | 60 | 30 | 240 | 3 x 10 sec; 2M MgCl2 | 50 |
| Fig 5C | Single Cycle | barr1 WT | 411 | V2Rpp | 40000 | | | 3 | 60 | 30 | 240 | 3 x 10 sec; 2M MgCl2 | 50 |
| Fig 5C | Single Cycle | barr1 WT | 411 | Fab30 | 1000 | PIP2 | 40000 | 3 | 60 | 30 | 240 | 3 x 10 sec; 2M MgCl2 | 50 |
| Fig 5C | Single Cycle | barr1 WT | 411 | Fab30 | 1000 | V2Rpp | 40000 | 3 | 60 | 30 | 240 | 3 x 10 sec; 2M MgCl2 | 50 |
| Fig 5C | Single Cycle | barr1 WT | 411 | Fab30 | 1000 | PI(3)P | 40000 | 3 | 60 | 30 | 240 | 3 x 10 sec; 2M MgCl2 | 50 |
| Fig 5C | Single Cycle | barr1 WT | 411 | Fab30 | 1000 | PG | 40000 | 3 | 60 | 30 | 240 | 3 x 10 sec; 2M MgCl2 | 50 |
| Fig 5C | Single Cycle | barr1 3Q | 392 | Fab30 | 1000 | | | 3 | 60 | 30 | 240 | 3 x 10 sec; 2M MgCl2 | 50 |
| Fig 5C | Single Cycle | barr1 3Q | 392 | PIP2 | 40000 | | | 3 | 60 | 30 | 240 | 3 x 10 sec; 2M MgCl2 | 50 |
| Fig 5C | Single Cycle | barr1 3Q | 392 | PI(3)P | 40000 | | | 3 | 60 | 30 | 240 | 3 x 10 sec; 2M MgCl2 | 50 |
| Fig 5C | Single Cycle | barr1 3Q | 392 | PG | 40000 | | | 3 | 60 | 30 | 240 | 3 x 10 sec; 2M MgCl2 | 50 |
| Fig 5C | Single Cycle | barr1 3Q | 392 | V2Rpp | 40000 | | | 3 | 60 | 30 | 240 | 3 x 10 sec; 2M MgCl2 | 50 |
| Fig 5C | Single Cycle | barr1 3Q | 392 | Fab30 | 1000 | PIP2 | 40000 | 3 | 60 | 30 | 240 | 3 x 10 sec; 2M MgCl2 | 50 |
| Fig 5C | Single Cycle | barr1 3Q | 392 | Fab30 | 1000 | V2Rpp | 40000 | 3 | 60 | 30 | 240 | 3 x 10 sec; 2M MgCl2 | 50 |
| Fig 5C | Single Cycle | barr1 3Q | 392 | Fab30 | 1000 | PI(3)P | 40000 | 3 | 60 | 30 | 240 | 3 x 10 sec; 2M MgCl2 | 50 |
| Fig 5C | Single Cycle | barr1 3Q | 392 | Fab30 | 1000 | PG | 40000 | 3 | 60 | 30 | 240 | 3 x 10 sec; 2M MgCl2 | 50 |
| Fig 5C | Single Cycle | barr1 1-382 | 424 | Fab30 | 1000 | | | 3 | 60 | 30 | 240 | 3 x 10 sec; 2M MgCl2 | 50 |
| Fig 5C | Single Cycle | barr1 1-382 | 424 | PIP2 | 40000 | | | 3 | 60 | 30 | 240 | 3 x 10 sec; 2M MgCl2 | 50 |
| Fig 5C | Single Cycle | barr1 1-382 | 424 | PI(3)P | 40000 | | | 3 | 60 | 30 | 240 | 3 x 10 sec; 2M MgCl2 | 50 |
| Fig 5C | Single Cycle | barr1 1-382 | 424 | PG | 40000 | | | 3 | 60 | 30 | 240 | 3 x 10 sec; 2M MgCl2 | 50 |
| Fig 5C | Single Cycle | barr1 1-382 | 424 | V2Rpp | 40000 | | | 3 | 60 | 30 | 240 | 3 x 10 sec; 2M MgCl2 | 50 |
| Fig 5C | Single Cycle | barr1 1-382 | 424 | Fab30 | 1000 | PIP2 | 40000 | 3 | 60 | 30 | 240 | 3 x 10 sec; 2M MgCl2 | 50 |
| Fig 5C | Single Cycle | barr1 1-382 | 424 | Fab30 | 1000 | V2Rpp | 40000 | 3 | 60 | 30 | 240 | 3 x 10 sec; 2M MgCl2 | 50 |
| Fig 5C | Single Cycle | barr1 1-382 | 424 | Fab30 | 1000 | PI(3)P | 40000 | 3 | 60 | 30 | 240 | 3 x 10 sec; 2M MgCl2 | 50 |
| Fig 5C | Single Cycle | barr1 1-382 | 424 | Fab30 | 1000 | PG | 40000 | 3 | 60 | 30 | 240 | 3 x 10 sec; 2M MgCl2 | 50 |

**Supplemantary Data Table 3**
